# Oligomannose Fc Glycans Reprogram the Energetic and Conformational Basis of CD16a Recognition

**DOI:** 10.64898/2026.08.07.738819

**Authors:** Natesan Mani, Alla Polozova, Srirupa Chakraborty

## Abstract

IgG1 Fc recognition by FcγRIIIa/CD16a is a central determinant of antibody-dependent cellular cytotoxicity and is strongly regulated by Fc N297 glycosylation. While afucosylation and galactosylation have been extensively studied, the structural basis by which oligomannosidic Fc glycans modulate CD16a binding remains less clear, despite their prevalence in therapeutic antibodies and association with accelerated serum clearance. Here, we use all-atom molecular dynamics simulations to investigate how mannose-5 (M5) Fc glycosylation alters IgG1 Fc–CD16a recognition across Paired Biantennary (complex glycans on both Fc), asymmetric Unpaired (complex glycan on one Fc arm and M5 on the other), and Paired M5 glycoforms. Computed interaction energies reproduce the experimental trend that Paired M5 glycoforms bind CD16a less favorably than complex-type paired glycans, supporting the use of the simulations to interrogate the structural origin of this energetic hierarchy. Residue-wise energetic decomposition and contact analyses show that Paired M5 glycosylation redistributes energetic contributions away from the productive Fc–CD16a interface and reduces both protein-mediated and glycan-mediated physical contacts. Free energy surface analyses further reveal that Paired M5 systems sample broader, less stable receptor-bound conformational ensembles, while dynamic cross-correlation analysis shows reduced intra-domain and inter-domain coupling across the complex. Importantly, a single M5 glycan is sufficient to perturb productive recognition by increasing Fc-arm separation heterogeneity, reducing high-frequency protein contacts, and weakening long-range dynamic communication. Glycan identity on the receptor-proximal Fc arm emerges as a decisive determinant of binding, indicating that Fc glycan composition, pairing, and receptor-bound placement jointly encode CD16a recognition. Together, these findings provide a mechanistic framework for understanding how oligomannose Fc glycans remodel antibody–receptor engagement and suggest that asymmetric Fc glycosylation, combined with residue-level interface engineering, may offer new strategies for tuning therapeutic antibody effector function.

## INTRODUCTION

The interaction between the Fc domain of immunoglobulin IgG1 Fc and Fcγ receptors is one of the central molecular determinants of antibody effector function. By engaging Fcγ receptors on immune cells, IgG1 antibodies trigger antibody-dependent cellular cytotoxicity (ADCC), phagocytosis, and other innate immune responses that underpin the efficacy of both natural and therapeutic antibodies^1, 2^. These effects are mediated by residues in the CH2 domain, where the FcγR directly contacts the antibody. A defining feature of this interaction is its sensitivity to glycosylation. Each arm of the IgG1 Fc carries a conserved N-glycosylation site at asparagine N297, and changes in glycan composition can dramatically alter Fcγ receptor binding and downstream immune activity^3, 4^. Understanding how individual glycan structures modulate Fcγ receptor recognition is therefore a major goal in the rational design of next-generation antibody therapeutics. These Fc glycans regulate binding strength to Fcγ receptors expressed on macrophages, natural killer (NK) cells, neutrophils, and other immune cells, thereby influencing effector responses, including ADCC and phagocytosis^4–9^. As illustrated in **Figure 1**, ADCC is initiated when pathogen-bound antibodies are recognized by FcγRIII receptors on NK cells, with the glycans at the Fc-FcγRIII interface serving as key modulators of this interaction. Controlling Fc glycosylation has consequently become a widely used strategy for tuning antibody activity. The N297 glycans on the Fc and the N45 and N162 glycans on FcγRIIIa (CD16a) participate directly in complex formation and strongly influence binding affinity^5, 9^. Studies over the past decade have established that even subtle changes in Fc glycan composition can profoundly alter receptor recognition, with core fucosylation representing one of the most influential determinants of FcγRIIIa binding. Removal of core fucose markedly increases receptor affinity and ADCC activity^5, 10–12^; while increasing levels of terminal galactosylation can further enhance FcγRIIIa engagement^10, 12, 13^.

**Figure 1.**
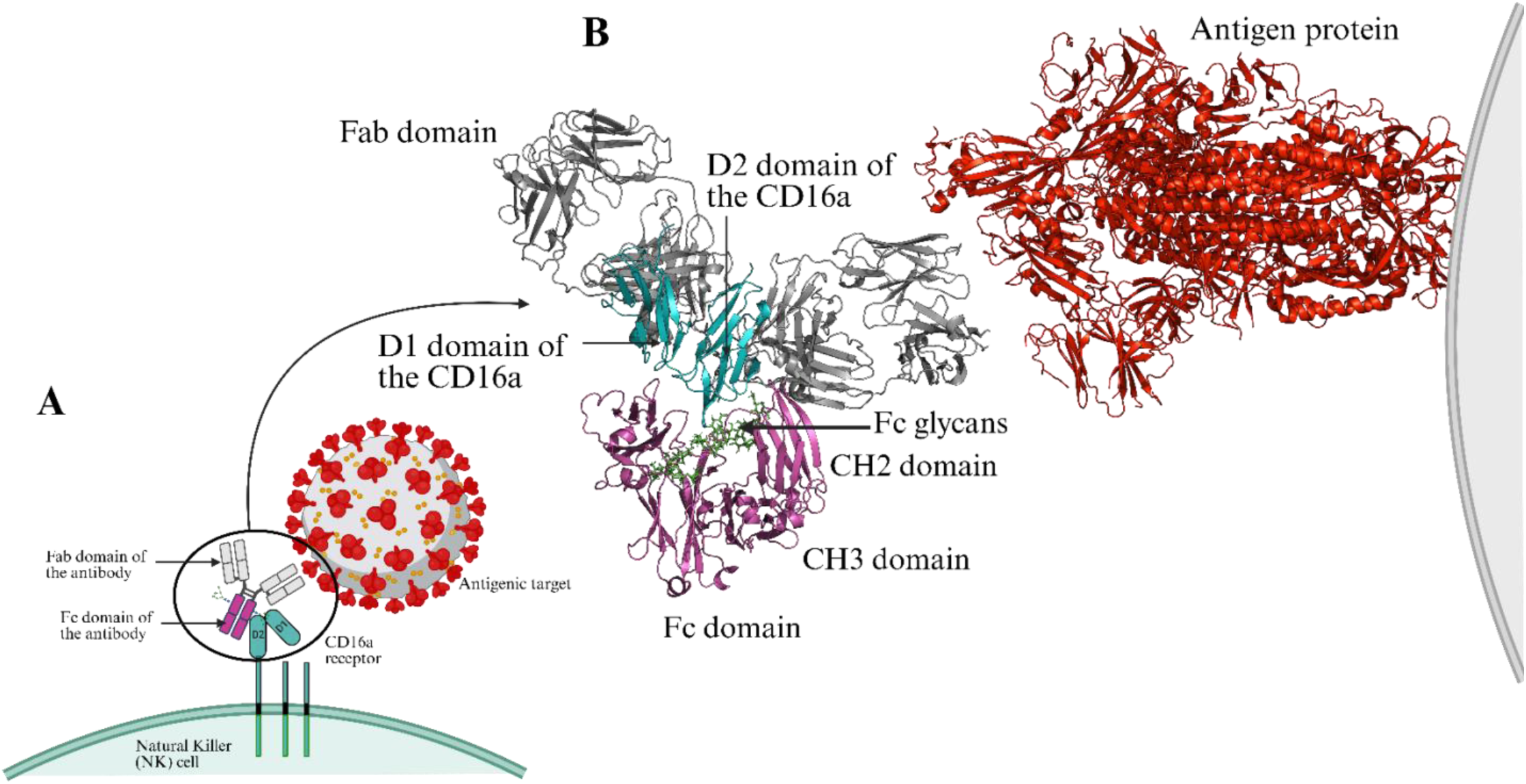
**Structural overview of antibody-mediated recognition during antibody-dependent cellular cytotoxicity (ADCC**). The IgG1 antibody acts as a bridge between the antigenic targets and the Natural Killer (NK) cell. The Fab domain binds the antigen displayed on the target cell surface, while the Fc domain is decorated with N-linked glycans at Asn297 and engages the CD16a (FcγRIII) receptor on the NK cell. **A**. Schematic illustration of this bridging interaction, showing the spatial arrangement of the antibody Fab and Fc domains, Fc glycans, and the CD16a receptor at the NK cell membrane. **B.** Molecular model of the ternary Fab–Fc–CD16a complex is shown. The Fc domain (magenta) bearing Fc glycans (green sticks) is contacted by the Fc-proximal D2 domain and Fc-distal D1 domain (teal) of CD16a, while the Fab domains (grey) project toward the antigen (red) on the opposing target cell surface.

High mannose glycans represent another important but comparatively less understood class of Fc glycoforms. Among these, five-mannose (M5) is one of the most prevalent high-mannose structures observed on therapeutic antibodies^14^. High mannose Fc glycans are associated with accelerated serum clearance^14, 15^, yet their impact on FcγRIIIa recognition remains incompletely understood. Experimental studies suggest that M5 glycosylation reduces FcγRIIIa binding relative to Paired Biantennary glycans^16^, but the structural basis for this behavior remains unclear. In contrast, on the receptor side, glycosylation effects are much better established. N45 and N162 are key determinants of Fc engagement^5, 9^: minimally processed glycans at these sites raise IgG1 Fc affinity^5, 17, 18^, and an M5 glycan at N162 increases affinity roughly 100-fold compared to a complex glycan^8, 17^. These observations highlight that glycan effects on antibody-receptor interactions are highly context-dependent and motivate a deeper mechanistic understanding of how Fc glycan composition influences receptor recognition. An additional layer of complexity arises from the intrinsic asymmetry of the FcγRIIIa–Fc interaction. Throughout this study, we designate the receptor-proximal Fc arm as Chain N (near) and the receptor-distal arm as Chain F (far) (see **Figure 2**). Recent experimental studies have shown that asymmetric glycosylation of the two Fc arms can significantly alter FcγRIIIa binding affinity^19^. Crystal structures and biochemical studies have shown that CD16a does not engage the two Fc chains equivalently^10, 20^. Chain N forms extensive contacts with the D2 domain of CD16a through the lower hinge region, including hydrogen bonds, salt bridges, and van der Waals interactions.

**Figure 2:**
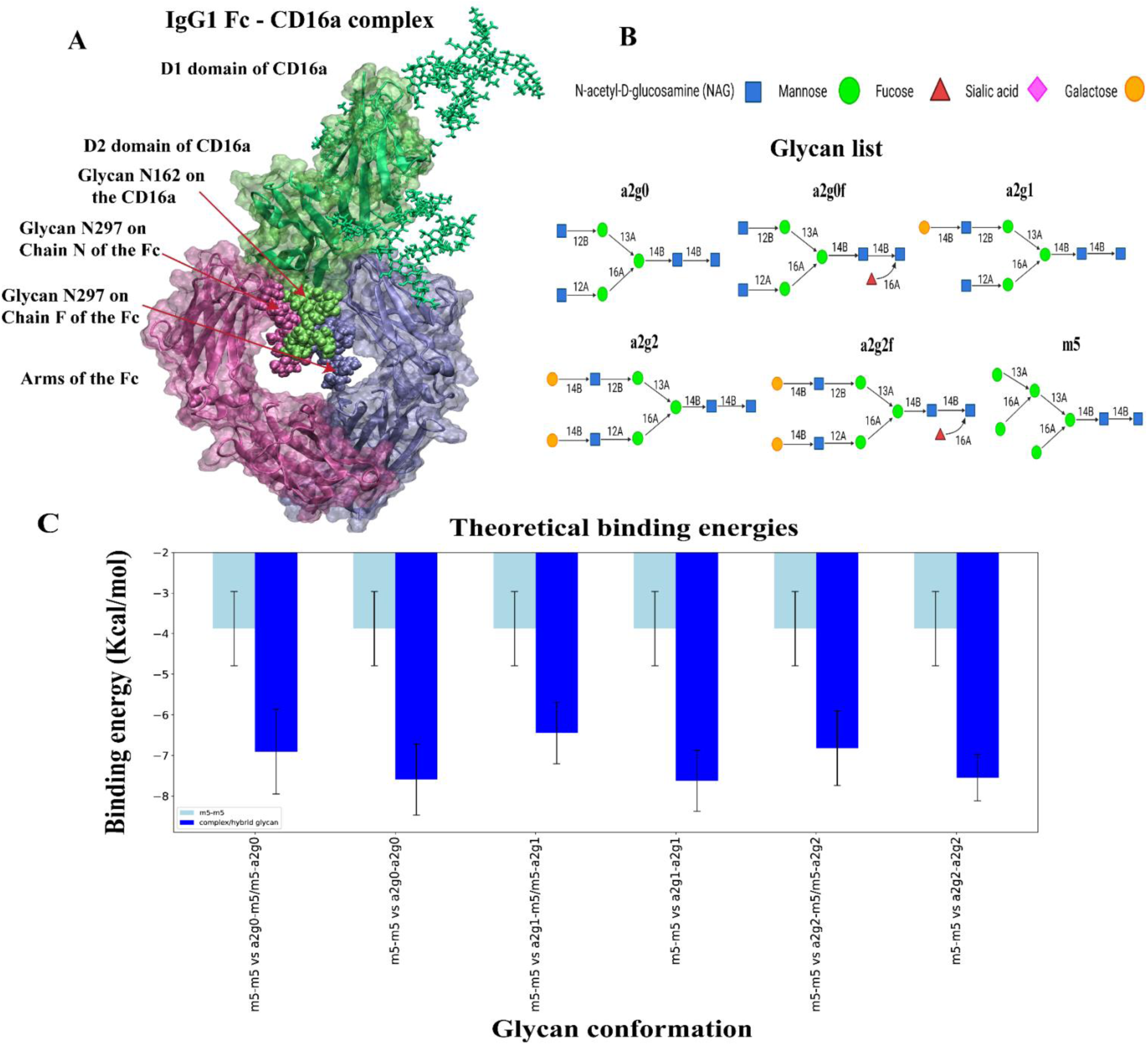
Paired high mannose glycosylation substantially reduces Fc-CD16a binding affinity. MMPBSA binding energy analysis quantifies the impact of Fc glycan composition on complex stability. **Panel A** depicts the IgG1 Fc-CD16a binding complex with the Fc domain (magenta and violet) and CD16a domain (lime). **Panel B** illustrates the N-glycan repertoire used in this study at Fc N297 and CD16a glycosylation sites. **Panel C** presents computed binding energies for afucosylated Paired biantennary, Unpaired, and Paired M5 glycosylated systems. Both Paired biantennary and Unpaired glycosylation confer substantially higher binding affinities compared to the Paired M5 variants, demonstrating that high mannose presence on both Fc chains dramatically destabilizes the Fc-CD16a interaction. These findings establish that Paired M5 glycosylation significantly disrupts the binding affinities of the IgG1 Fc-CD16a complex

Despite these advances, the molecular basis by which oligomannosidic Fc glycans regulate FcγRIIIa recognition remains unresolved. Most structural and mechanistic work has focused on receptor glycosylation, core fucosylation, or terminal galactosylation, whereas the effect of M5 glycans on the Fc itself has been primarily described through clearance and binding measurements. As a result, it remains unclear whether M5 glycosylation on the Fc weakens receptor engagement through local loss of interfacial contacts, broader changes in the Fc–CD16a conformational ensemble or altered dynamic coupling across the bound complex. Moreover, because CD16a engages the two Fc arms asymmetrically, it is not known whether M5 glycans on the receptor-proximal and receptor-distal Fc arms are functionally equivalent. These observations raise a central mechanistic question: does oligomannosidic M5 glycosylation weaken FcγRIIIa recognition solely because of its glycan composition, or is its effect amplified by the asymmetric geometry of the Fc–CD16a interface? More specifically, is a single M5 glycan sufficient to perturb receptor engagement, and does M5 on the receptor-proximal Fc arm disrupt binding more strongly than M5 on the distal arm? Answering these questions is essential for understanding whether FcγRIIIa binding is governed only by the chemical identity of the Fc glycans, or by a coupled dependence on glycan identity, glycan placement, and the conformational dynamics they induce within the bound complex.

To define the molecular mechanism underlying this composition- and position-dependent glycan effect, we performed extensive all-atom Molecular Dynamics (MD) simulations of the IgG1 Fc–CD16a complex across a panel of glycosylation states (see **SI Table 1**). We focused on CD16a because this Fcγ receptor binds IgG1 with moderate affinity yet strongly mediates ADCC, and both properties can be tuned by modifying the glycoforms of the Fc and CD16a^21, 22^. Our study encompasses the following: (i) ‘Paired Biantennary’ systems containing complex biantennary glycans on both Fc arms, (ii) ‘Unpaired’ systems contain a biantennary complex glycan on one arm and an oligomannose M5 glycan on the other, and (iii) ‘Paired M5’ systems contain oligomannose glycans on both Fc arms (refer **SI Table 1**). Three independent 1.2 μs simulations were performed for each system (with 13 systems in total). These timescales provided sufficient sampling to allow glycosylation-dependent changes in binding energetics, protein-protein interactions, glycan-mediated contacts, conformational free-energy landscapes, and correlated domain motions. Through these analyses, we investigated how oligomannose glycans influence FcγRIIIa binding, whether a single oligomannose glycan is sufficient to perturb receptor engagement, and how glycan positioning within the asymmetric Fc–CD16a interface shapes these effects. Our results show that oligomannosidic Fc glycosylation weakens Fc–CD16a recognition not just by changing glycan composition, but by disrupting productive interfacial contacts, destabilizing favorable receptor-bound conformations, and suppressing long-range dynamic coupling across the asymmetric Fc–CD16a interface. This disruption is strongly position dependent: M5 on the receptor-proximal Fc arm produces the greatest impairment, whereas a complex glycan at this position preserves more favorable receptor engagement. Together, these findings suggest a broader principle: FcγRIIIa binding is governed not only by glycan composition but also by glycan placement within an inherently asymmetric binding interface, providing new mechanistic insights and design rules for glycoengineering therapeutic antibodies with tunable effector functions.

## RESULTS

### Glycoform Design and Simulation Overview to Compare Mannose and Complex-type Fc Glycans

To determine how high-mannose Fc glycans alter IgG1 Fc–CD16a recognition relative to complex-type Fc glycans, we performed all-atom MD simulations on a panel of 13 IgG1 glycosylation patterns (**SI Table 1**). These systems were designed to separate three related effects: the impact of paired high-mannose M5 glycans, the effect of introducing a single M5 glycan on one Fc arm, and the influence of glycan position relative to the bound CD16a receptor. All systems were constructed from the crystal structure of the Fc-CD16a complex (PDB code: 1E4K) in RCSB^20^. CD16a glycosylation was generally modelled using the most probable glycan at each site from mass-spectrometry data^8^, but the N162 was modelled as a high-mannose glycan because it elicits the strongest binding to the IgG1 Fc^5^. The simulations are grouped into three categories: (i) Paired Biantennary, with the same complex glycan on both Fc arms; (ii) Unpaired, with a complex glycan on one arm and a high-mannose (M5) glycan on the other; and (iii) Paired M5 oligomannose, with 5-mannose on both arms. The image of the complex with the different glycan types used in this study is shown in **Figures 2A** and **2B**. The Paired Biantennary and Unpaired glycans were varied with increasing degrees of galactosylation from 0 to 2 and are denoted G0, G1, and G2 for zero, one, and two galactose residues per complex glycan antenna, respectively. As noted above, to account for asymmetric CD16a binding, Chain N denotes the Fc arm that is proximal to the receptor and Chain F, the distal arm. For Unpaired systems, the notation a2gx-m5 (x = 0, 1, 2) indicates that the complex glycan occupies the proximal arm and the mannose the distal arm; the reverse notation applies for the opposite arrangement. All the results for these Unpaired systems are from the combined simulations of these pairs, and the binding energies reported for each Unpaired system represent a simple (50:50) average over the two glycan-position combinations (complex glycan on the Fc proximal versus distal positioning. **Figure 2** and **SI Table 1** illustrate the different glycans modelled on the Fc. To highlight the mechanistic differences between Paired Biantennary, and Paired M5 systems, we selected the G1 type of Paired/Unpaired systems of each for comparison in the main manuscript. Results for the mechanistic comparison for the other Paired/Unpaired systems (G0 and G2) against Paired M5 are provided in the SI. The final list of systems is represented in **SI Table 1**. Three independent 1.2 µs runs were performed in AMBER using the CHARMM36m force field^23^. The first 200 ns of each trajectory were discarded to ensure convergence before analysis. Details of system setup and simulation protocol can be found in the **Methods** section. This simulation design enabled direct comparison of how Fc glycan composition, pairing, and receptor-proximal positioning affect Fc–CD16a binding energetics, interfacial contacts, glycan organization, and protein dynamics.

### Oligomannosidic M5 glycosylation on Fc weakens CD16a binding in a pairing and position-dependent manner

Mannose-5 (M5) is the predominant high-mannose form observed in therapeutic antibodies, and high-mannose Fc glycosylation has been associated with increased antibody clearance relative to other glycan types^14–16^. However, the structural and energetic consequences of M5 glycosylation for Fc–CD16a complex formation remain incompletely understood. Recent experimental measurements suggest that Paired M5 Fc glycosylation has lower CD16a binding affinity than Paired Biantennary or Unpaired glycosylation, although the molecular basis for this effect remains unclear^16^. To assess relative interaction strengths across the simulated systems and quantify the impact of Paired M5 glycosylation on IgG1 Fc–CD16a complex formation, we computed an MM/PBSA-style interaction energy estimate from the MD trajectories using AmberTools-based energetic decomposition^24, 25^. The Van der Waals and Electrostatic interaction terms were empirically scaled (α = 0.158 and β = 0.153, respectively; see **Methods**), following the approach of Li et al., to place the results on a scale more comparable to experimental readouts^26^. Importantly, these calculations were not intended to provide absolute binding free energies, but rather to quantify comparative energetic changes associated with glycan composition. We first compared Paired M5 glycosylation against the high-affinity afucosylated Paired Biantennary and Unpaired systems across different degrees of galactosylation (**Figure 2C**). Paired Biantennary and Unpaired systems showed substantially stronger Fc–CD16a interaction energies than Paired M5 systems. This indicates that placing M5 glycans on both Fc arms markedly weakens CD16a engagement, consistent with experimental observations that Paired M5 glycosylation reduces receptor binding relative to complex-type Fc glycosylation^16^.

We next asked whether this energetic penalty requires M5 on both Fc arms and whether a single M5 glycan is sufficient to perturb receptor engagement. To address this, we compared Paired Biantennary systems with their corresponding Unpaired counterparts across G0, G1, and G2 galactosylation states. Across all three galactosylation levels, Paired Biantennary systems showed stronger computed binding energies than their Unpaired counterparts (**Figure 3A, Blocks 1–3**). Thus, even a single M5 substitution reduces Fc–CD16a binding relative to the corresponding paired complex-type glycoform.

**Figure 3:**
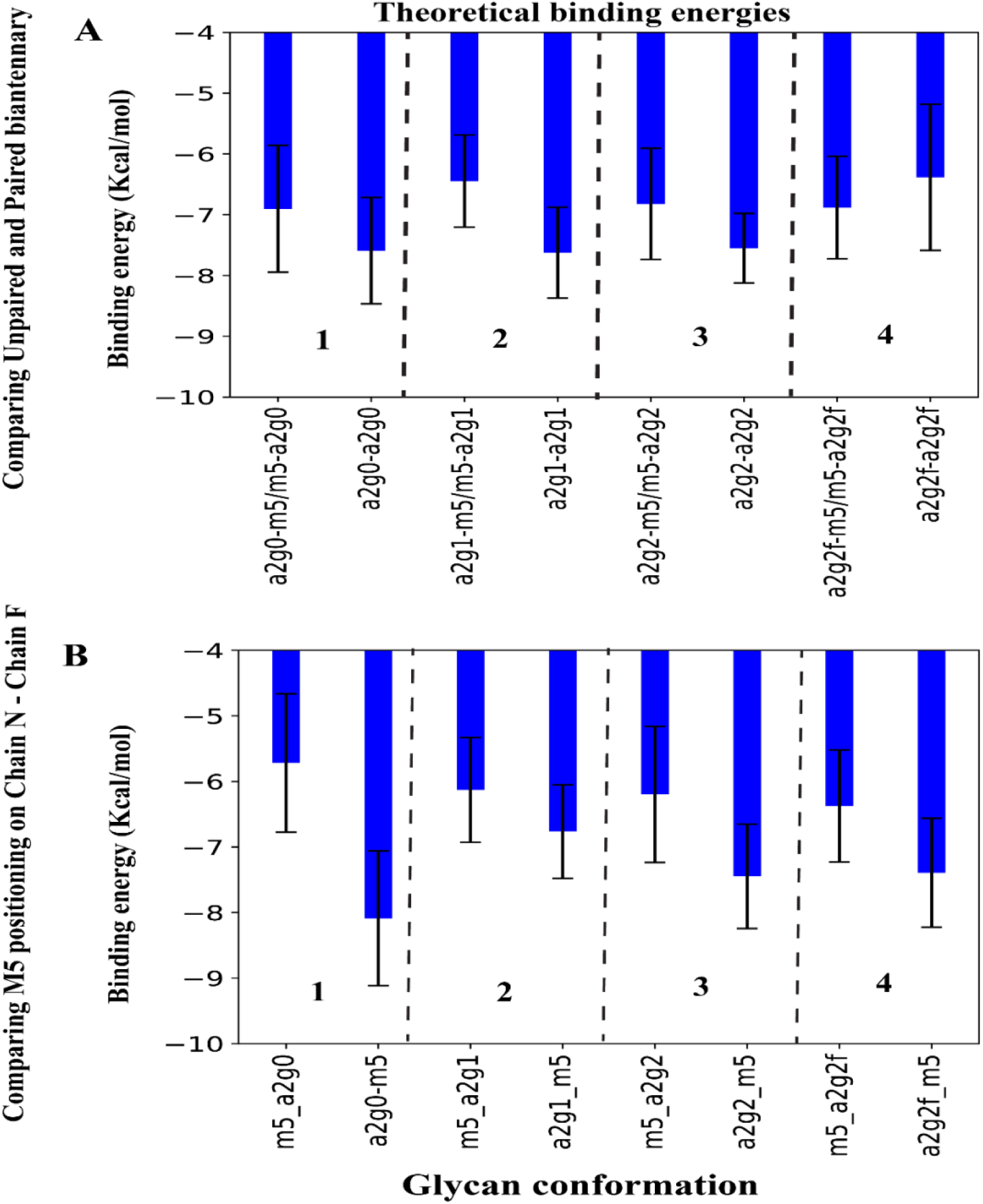
The presence and position of a single high mannose dictate the affinity of the Fc-CD16a binding complex. We compared the theoretical binding energy values for high-affinity (afucosylated) Unpaired and Paired biantennary glycosylated systems (see **Panel A**), with the difference between the two being the presence or absence of high mannose. We find that Paired biantennary systems have a higher binding affinity than Unpaired, with the only exception being **Block 4**, which has also been identified in literature. In **Panel B,** we compare different systems regarding the position of the High mannose on the Fc: either on the proximal arm to CD16a (on the left of each block) or on the distal arm (on the right of each block). We observe in almost all cases that the presence of High mannose on the proximal arm leads to a reduction in binding affinity values, emphasizing that both the presence and position of High mannose have an immense impact on the formation of the IgG1 Fc-CD16a binding complex.

Because fucosylation can alter the broader Fc glycan architecture and is known to modulate FcγRIIIa recognition, we also examined fucosylated variants of the highest-affinity systems. Fucose was added to the G2 Paired Biantennary system, yielding ‘a2g2f–a2g2f’, and to the corresponding Unpaired systems, yielding ‘a2g2f–m5’ and ‘m5–a2g2f’, as shown in **Figure 3A, Block 4.** We find that the Unpaired system had a higher binding affinity as compared to the Paired Biantennary system with the presence of fucose, and a similar trend was also captured from experiments by Meudt et al.^16^. This suggests that the effect of a single M5 glycan is modulated by the broader Fc glycan context, including the presence of fucose. Our structural findings were able to explain these calculated and experimentally observed binding affinity differences as discussed in subsequent results (**Figure 10**).

Finally, because CD16a binds Fc asymmetrically, we examined whether the position of the single M5 glycan influences binding. In the Unpaired systems, M5 on Chain N, the receptor-proximal Fc arm, reduced binding relative to the corresponding system in which the complex glycan occupied Chain N (**Figure 3B, Blocks 1–4**). The same positional trend was observed for the fucosylated pair a2g2f–m5 and m5–a2g2f (**Figure 3B, Block 4**), indicating that even in higher-affinity fucosylated systems, productive receptor engagement is favored when the complex-type glycan is located on the CD16a-proximal Chain N. This observation is further explored in a later section. Together, these energetic comparisons show that oligomannosidic Fc glycosylation weakens Fc–CD16a engagement in a pairing- and position-dependent manner. Paired M5 produces the strongest loss in interaction strength, while a single M5 glycan is also sufficient to reduce binding, particularly when positioned on the receptor-proximal Fc arm.

### Paired M5 glycosylation redistributes interfacial energetics and reduces inter-protein and inter-glycan contacts

The weaker binding energies observed for Paired M5 systems suggested that oligomannosidic Fc glycosylation disrupts CD16a engagement. We next asked whether this reduction reflects a uniform weakening of the Fc–CD16a interface or a reorganization of the energetic and physical interactions that stabilize the bound complex. To address this, we performed residue-wise energetic decomposition using the MMPBSA.py module of AMBER^24, 25^ and identified the top 20 energetically active Fc (top 5% of Fc residues) and CD16a (top 10% of CD16a residues) residues that contributed most strongly to binding within the high-affinity Paired Biantennary/Unpaired systems and within the Paired M5 systems.

Mapping these residues onto representative Fc–CD16a structures revealed a clear spatial reorganization of the energetic landscape (see **Figure 4** for G1 systems and **SI Figure 2** for the other glycoforms). In the Paired Biantennary and Unpaired systems, energetically important residues were concentrated near the Fc–CD16a binding interface, consistent with productive receptor engagement. By contrast, in the Paired M5 systems, several energetically active CD16a residues were located farther away from the interface and redistributed toward regions of CD16a that do not form the primary Fc-contacting surface, including parts of the D2 domain (**Figure 4B, D**). Thus, Paired M5 glycosylation does not simply reduce the magnitude of Fc–CD16a binding energy; it redirects energetic contributions away from the productive receptor-binding interface. The top 10 contributing residues for the Unpaired G1, Paired G1, and Paired M5 systems are listed in **SI Table 2**. Because residues were ranked by their relative contribution within each system, these sets are not mutually exclusive, meaning that in the lower-affinity Paired M5 systems, residues that fall below the top-contribution cutoff in the higher-affinity Paired Biantennary/Unpaired systems can rank among the leading contributors. To determine whether this energetic redistribution was accompanied by a physical loss of contacts, we next quantified three classes of interactions: (i) inter-protein contacts between Fc and CD16a, (ii) inter-Fc glycan contacts between the two N297 glycans, and (iii) Fc–CD16a glycan contacts between the Fc N297 glycans and the N162 glycan on CD16a. Difference contact maps (see **Figure 5** for G1 and **SI Figure 3** for the other glycoforms) comparing Paired Biantennary/Unpaired systems against Paired M5 systems showed that the Paired Biantennary and Unpaired systems dominate the Fc–CD16a protein–protein contact landscape, whereas Paired M5 systems lose many of the high-frequency contacts that characterize the productive receptor-bound interface (**Figure 5A, 5B** and **SI Figure 3A–D**). Several residues identified as energetically important in the decomposition analysis also appeared within these contact regions and are marked on the contact maps (**Figure 5A, 5B**), supporting a direct connection between the energetic redistribution observed in **Figure 4** and the physical weakening of the Fc–CD16a interface.

**Figure 4:**
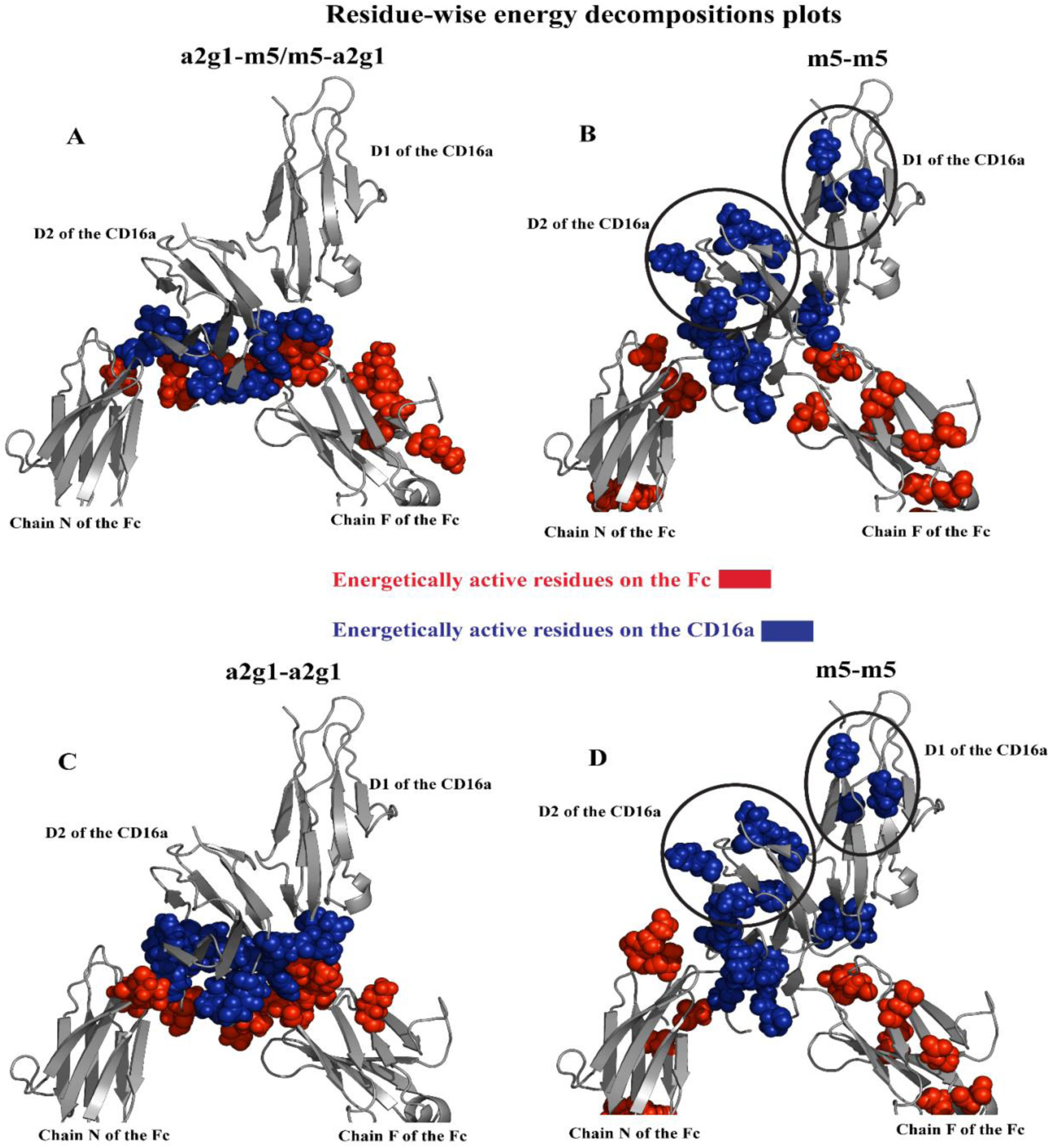
Paired M5 glycosylation redistributes the Fc–CD16a binding interface. Energy decomposition analysis showing the 20 most energetically active residues at the Fc–CD16a interface that are unique to each set of Paired/Unpaired and Paired M5 systems. **(A, C)** Unpaired and Paired biantennary glycosylated systems and **(B, D**) Paired M5 systems. Energetically critical CD16a residues in Paired M5 complexes (circled) are shifted away from the binding interface relative to Paired biantennary and Unpaired systems. These results mechanistically explain the reduced binding affinities observed in Figure 2. Refer to **SI Table 2** for the list of residues

**Figure 5:**
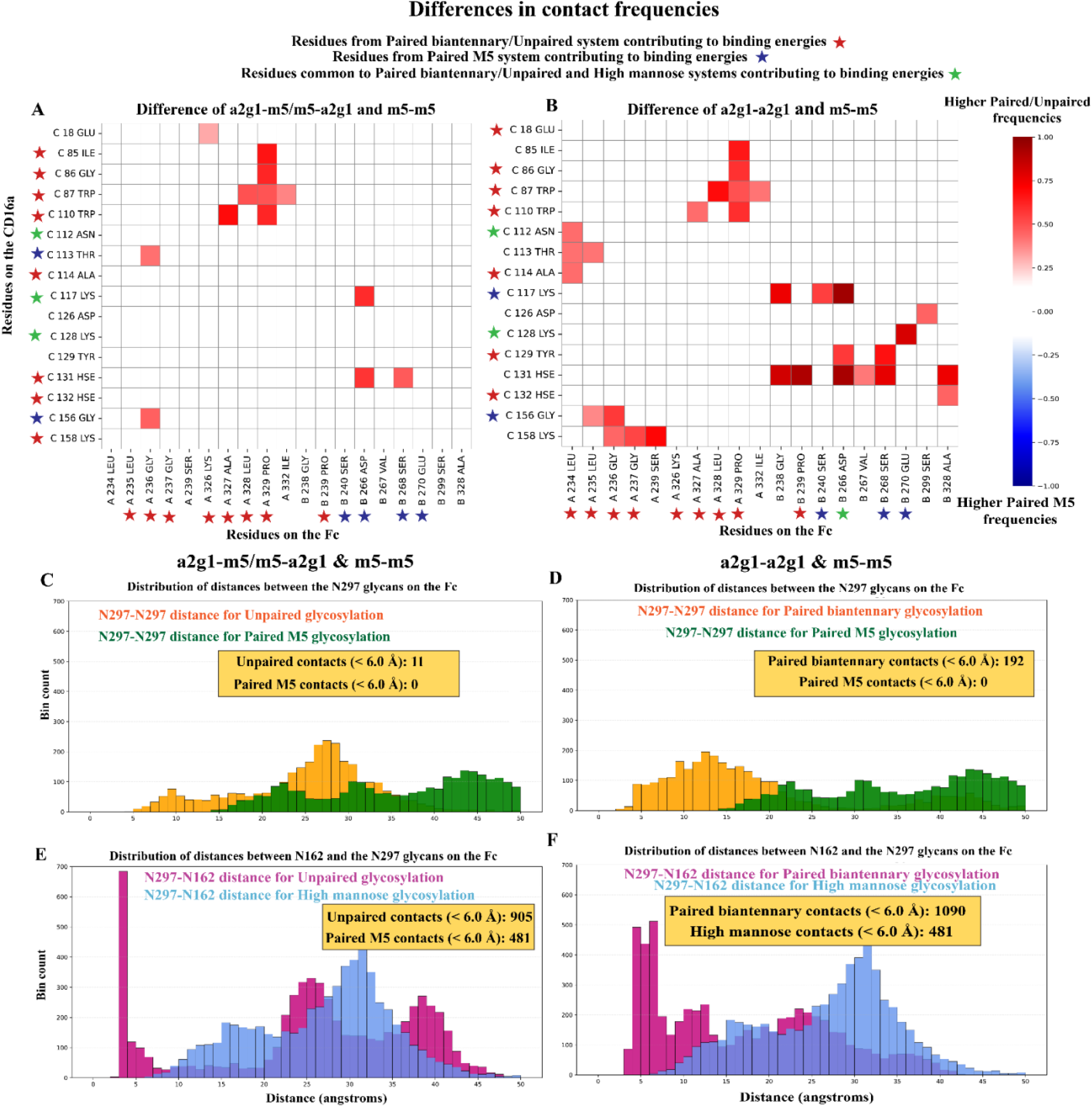
**Paired M5 glycosylation reduces protein–protein and glycan–glycan contacts at the Fc–CD16a interface**. Contact frequency analysis comparing Unpaired **(A, C, E)** and Paired biantennary **(B, D, F)** glycosylated systems against Paired M5 variants. **Difference in** protein–protein contacts – (**A**) Unpaired minus Paired M5; (**B**) Paired biantennary minus Paired M5**. (C, D)** Inter-glycan contacts at Fc N297. **(E, F**) Inter-glycan contacts between Fc N297 and CD16a N162, which aid complex formation. Unpaired and Paired biantennary systems show higher contact frequencies than Paired M5 variants across all three categories.

We then examined whether Paired M5 glycosylation also disrupts glycan-mediated contacts within the complex. Inter-Fc glycan contacts were quantified from heavy-atom distances between the two Fc N297 glycans, whereas Fc–CD16a glycan contacts were quantified from distances between the Fc N297 glycans and the CD16a N162 glycan, using a cutoff of 6.0 Å. Consistent with the protein–protein contact analysis, Paired Biantennary and Unpaired systems showed substantially greater glycan contact frequencies than Paired M5 systems (**Figure 5C–F** and **SI Figure 3E–H**). The two M5 glycans in the Paired M5 systems remain farther apart than Fc glycan pairs in systems where at least one Fc arm carried a complex-type glycan. In addition, Fc–CD16a inter-glycan contacts are significantly more frequent than inter-Fc glycan contacts across the systems, highlighting the dominant role of receptor-associated glycan engagement in organizing the Fc–CD16a interface. Thus, having Paired-M5 glycoform not only changes glycan interactions but extends to reorganization of the protein interface as well, that leads to a notable loss of productive engagement between Fc–CD16a. Together, these analyses show that Paired M5 glycosylation disrupts both the energetic and physical architecture of the Fc–CD16a interface. The shift of energetic contributors away from the productive binding surface, combined with the loss of several protein-mediated and glycan-mediated contacts, provides a structural explanation for the weaker interaction energies observed for Paired M5 systems.

### Paired M5 glycosylation destabilizes the binding complex and the conformational landscape

Having shown that Paired M5 glycosylation reduces binding energies and disrupts productive physical contacts at the binding interface, we next asked whether these local perturbations are accompanied by broader changes in the receptor-bound conformational ensemble. We tracked CD16a motion along two principal rotational axes together with the ‘opening’ of the Fc arms relative to CD16a (see **Methods**; **Figure 6G, 6H**; **and SI Figure 4K, 4L**). These collective coordinates were projected onto 2D free energy surfaces (FES), where deeper basins correspond to more frequently sampled and higher-stability conformational states. The FESs revealed a clear glycoform-dependent difference in conformational sampling. The Paired Biantennary and Unpaired systems sampled compact, well-defined basins, consistent with stable receptor-bound conformations (refer **Figure 6A–F** for G1 systems**, SI Figure 4A-J** for other glycoforms). These observations are consistent for G1 type of Unpaired/Paired glycans (**Figure 6A–F)** and also for G0, G2 types (**SI Figure 4A-J**). In contrast, the Paired M5 systems sampled broader and more diffuse basins across both CD16a rotational coordinates and Fc-opening coordinates, indicating a shift toward more heterogeneous and lower-stability receptor-bound ensembles. Thus, Paired M5 glycosylation not only reduces local interfacial contacts; it also broadens the conformational space accessible to the Fc–CD16a complex and destabilizes conformations associated with productive receptor engagement.

**Figure 6:**
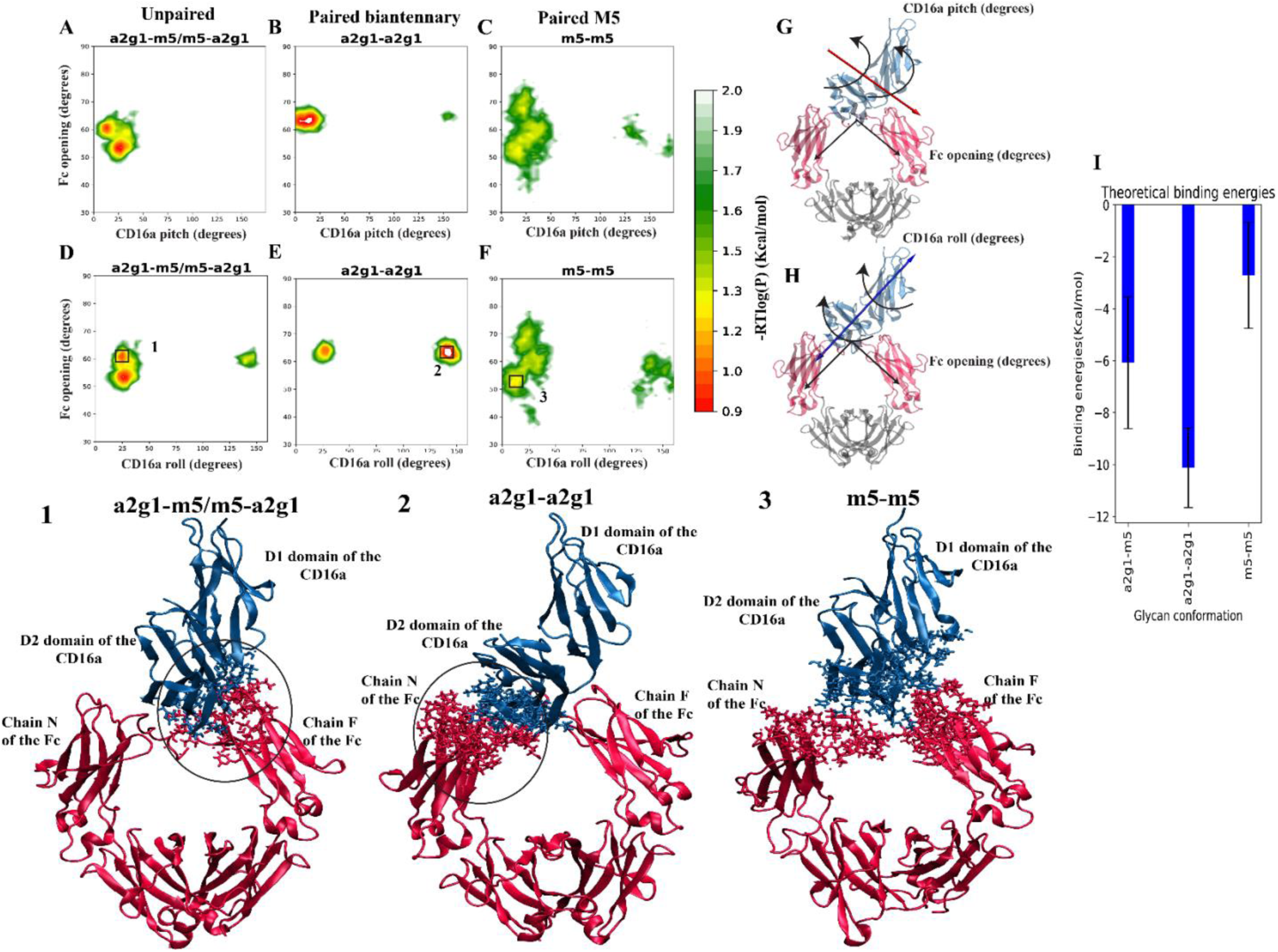
Paired M5 glycosylation destabilizes CD16a conformations. Free energy surface (FES) analysis of CD16a rotational motion along two independent axes; lower colorbar values indicate higher stability. **(A, B)** FES for Unpaired and Paired biantennary systems. **(C, D)** FES for Paired M5 variants, with extracted structures from stable basins labelled 1–3 (Unpaired, Paired biantennary, and Paired M5, respectively). For each system, the structure with the lowest RMSD from the trajectory corresponding to that basin is taken. The licorice sticks represent the atoms that are within 10 Å of the D2 domain and its proximal Fc domain. For the structure corresponding to a2g1-m5/m5-a2g1 (**Structure 1**, corresponding to basin 1 from the **Unpaired** plot of **Panel D**), we find the D2 domain oriented towards the CD16a distal arm of the Fc (Chain F) and for the structure corresponding to a2g1-a2g1 (**Structure 2**, corresponding to basin 2 from **Paired** plot of **Panel E**), we find that the D2 domain orients towards the CD16a proximal arm of the Fc (Chain N). In the Paired M5 structure (**Structure 3**, corresponding to basin 3 from the **Paired M5** plot of **Panel F**) the D2 domain of CD16a adopts an orientation that positions it away from both arms of the Fc region. The MMPBSA binding energies computed from the respective basins (**Panel I**), are in agreement with these mechanistic findings. (**E, F**) Rotational axes used for FES analysis. I

To connect these ensemble-level differences to specific receptor orientations, we extracted representative structures (RMSD closest to the mean) from the stable basins marked as regions **1–3** in the FES landscapes (**Figure 6D, E, F; Blocks 1–3**). The CD16a rotational coordinate shown in **Figure 6H** captures a rolling-type motion of the receptor across the Fc surface and reveals two stable basins for the Paired Biantennary and Unpaired systems, corresponding to receptor orientations closer to one or the other Fc arm. In the Paired Biantennary system, these two basins were nearly equally populated, indicating that the complex can stably sample both receptor orientations. In contrast, the Unpaired system showed a stronger population bias toward the basin in which CD16a is positioned closer to Chain F, the receptor-distal Fc arm in the crystal structure. Representative structures from these two basins, extracted at comparable Fc-arm opening angles, are shown as Structures **1** and **2**. In Structure **1**, corresponding to the Unpaired system, the CD16a D2 domain is oriented toward Chain F, whereas in Structure 2, corresponding to the Paired Biantennary system, we are representing the other FES basin, where the D2 domain is oriented toward Chain N, the receptor-proximal arm. Paired M5 samples a more heterogeneous binding ensemble, with CD16a sampling a broad range of placements between the two Fc arms and a larger variation in Fc-arm opening – with a larger propensity to sample around chain F (structure **3**). These representative structures recapitulated the energetic trends from the full trajectories, with the Paired Biantennary basin structure showing stronger interaction energy than the corresponding unbiased trajectory average for the same system (**Figure 6I; compare with Figure 3A, Block 2** for **a2g1–a2g1**). These results suggest that receptor orientations maintaining D2 proximity to Chain N are especially favorable for Paired Biantennary systems, whereas the presence of M5 shifts the complex toward less productive and more heterogeneous binding geometries.

### Paired M5 glycosylation suppresses long-range dynamic coupling across the Fc–CD16a complex

The FES analysis showed that Paired M5 glycosylation destabilizes the receptor-bound conformational ensemble. We next asked whether this conformational destabilization is accompanied by changes in long-range correlated motions across the Fc–CD16a complex. To address this, we computed dynamic cross-correlation matrices (DCCMs) using MD-TASK^27^. DCCM analysis quantifies time-correlated motions between residue pairs over the MD trajectories and provides a system-wide view of how motions within Fc, within CD16a, and across the Fc–CD16a interface are dynamically coupled.

To isolate the effect of Paired M5 glycosylation, we plotted DCCM difference maps between the higher-affinity Paired Biantennary/Unpaired systems and the Paired M5 systems (refer **Figure 7A,7B** for G1 and **SI Figure 5A–D** for the other glycoforms). The Paired Biantennary and Unpaired systems showed stronger intra-Fc and intra-CD16a correlations than the Paired M5 systems, as highlighted by the black boxes in **Figure 7A, 7B**. These systems also showed stronger inter-domain correlations between Fc and CD16a, highlighted by the green boxes, indicating more coordinated motion across the receptor-bound interface.

**Figure 7:**
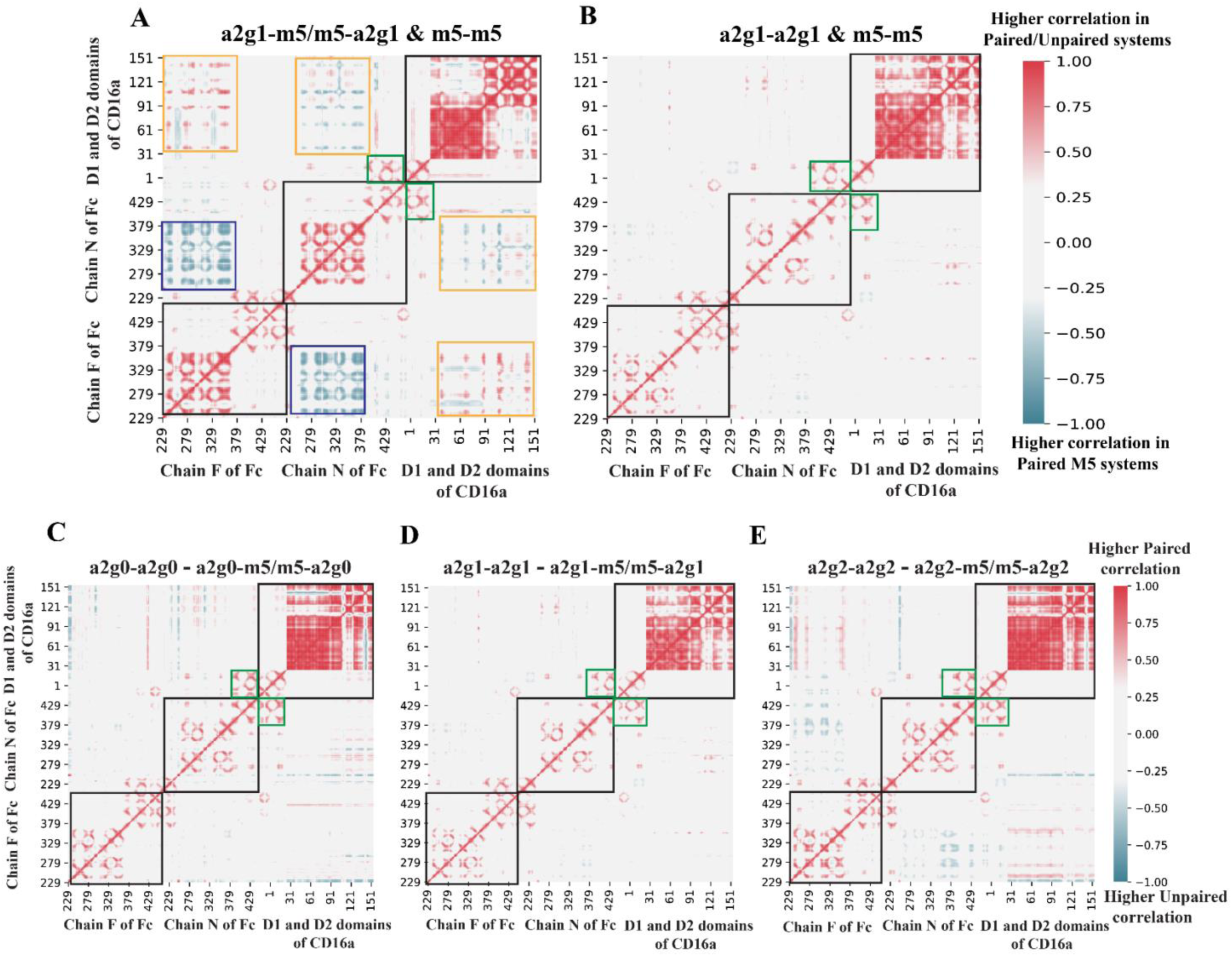
**Paired M5 glycosylation suppresses correlated domain motions in the Fc–CD16a complex**. Dynamic cross-correlation analysis across Paired biantennary, Unpaired, and Paired M5 glycosylated systems. (A, B) Difference plot of correlated motions in Paired biantennary and Unpaired systems versus Paired M5, highlighted are higher intra-Fc (black squares), inter Fc–CD16a (green squares), intra-CD16a (black squares) in the Paired/Unpaired systems, and elevated inter-Fc correlations in Paired M5 relative to Unpaired (blue squares, A). Paired biantennary/Unpaired systems show a higher degree of beneficial correlated motions relative to the Paired M5 systems. (C, D, E) Difference plot of correlated motions in Paired biantennary and Unpaired systems across varying galactose content. Paired biantennary systems maintain stronger correlated motions than Unpaired variants across all domains, independent of galactosylation level.

In contrast, Paired M5 glycosylation reduced these favorable intra-domain and inter-domain correlations. This loss of correlated motion is consistent with the broader, less defined conformational basins observed in the FES analysis and suggests that Paired M5 disrupts not only local interfacial contacts but also the larger-scale dynamic communication that stabilizes productive Fc–CD16a engagement. Compared to Unpaired, the Paired M5 system showed increased inter-Fc correlations in the blue-boxed region of **Figure 7A**, suggesting that M5 can promote Fc-arm motions that are not coupled productively to CD16a binding. Together, these results indicate that Paired M5 glycosylation suppresses the long-range dynamic coupling associated with stable Fc–CD16a recognition. Thus, the reduced binding energies and diminished physical contacts observed for Paired M5 systems are accompanied by a loss of coordinated motions across both the individual domains and the Fc–CD16a interface.

### A single M5 glycan is enough to significantly reduce dynamic coupling and physical contacts relative to Paired Biantennary glycosylation

The preceding analyses showed that Paired M5 glycosylation disrupts Fc–CD16a binding energetics, physical contacts, conformational stability, and long-range correlated motions. We next asked whether these effects require M5 on both Fc arms or whether a single M5 glycan is sufficient to perturb receptor engagement. This comparison is particularly important because the binding- energy analysis showed that high-affinity Unpaired systems, which contain one complex-type glycan and one M5 glycan, have lower computed binding affinities than their corresponding Paired Biantennary systems (**Figure 3A**). We went on to compare large-scale correlated motions between Paired Biantennary and Unpaired systems using DCCM difference maps (**Figure 7C–E**). Paired Biantennary systems showed stronger intra-Fc correlations, highlighted by the black boxes, and stronger inter-Fc–CD16a correlations, highlighted by the green boxes, than their Unpaired counterparts. These differences indicate that replacing one complex-type Fc glycan with M5 is sufficient to reduce both intra-domain coupling within Fc and long-range dynamic coupling across the Fc–CD16a interface.

We then examined whether this reduced dynamic coupling due to a single M5 was reflected in the conformational ensemble of the complex. Following the same FES-based analysis used above for the Paired M5 comparison, we projected CD16a rotational motions together with an Fc conformational coordinate onto two-dimensional free energy surfaces (**Figure 8**). For the Paired Biantennary versus Unpaired comparison, the Fc-opening coordinate is replaced by the distance between the centers of mass of the two Fc CH2 domains (**Figure 8G, H**). Paired Biantennary systems sampled more compact conformational spaces with deeper, higher-stability basins, whereas Unpaired systems sampled broader and more diffuse lower-stability conformational landscapes (**Figure 8A–F**). This difference was also reflected in the Fc-arm separation coordinate: the Paired Biantennary systems maintained a relatively uniform inter-Fc distance, whereas the Unpaired systems showed substantially greater variation in the distance between the two Fc arms. Thus, the presence of a single M5 glycan increases conformational heterogeneity not only in receptor orientation but also in the relative positioning of the Fc arms. These results are consistent with the DCCM analysis: stronger intra-Fc and inter-domain correlations in the Paired Biantennary systems correspond to more stable receptor-bound conformational ensembles, whereas reduced coupling in the Unpaired systems corresponds to increased conformational heterogeneity.

**Figure 8:**
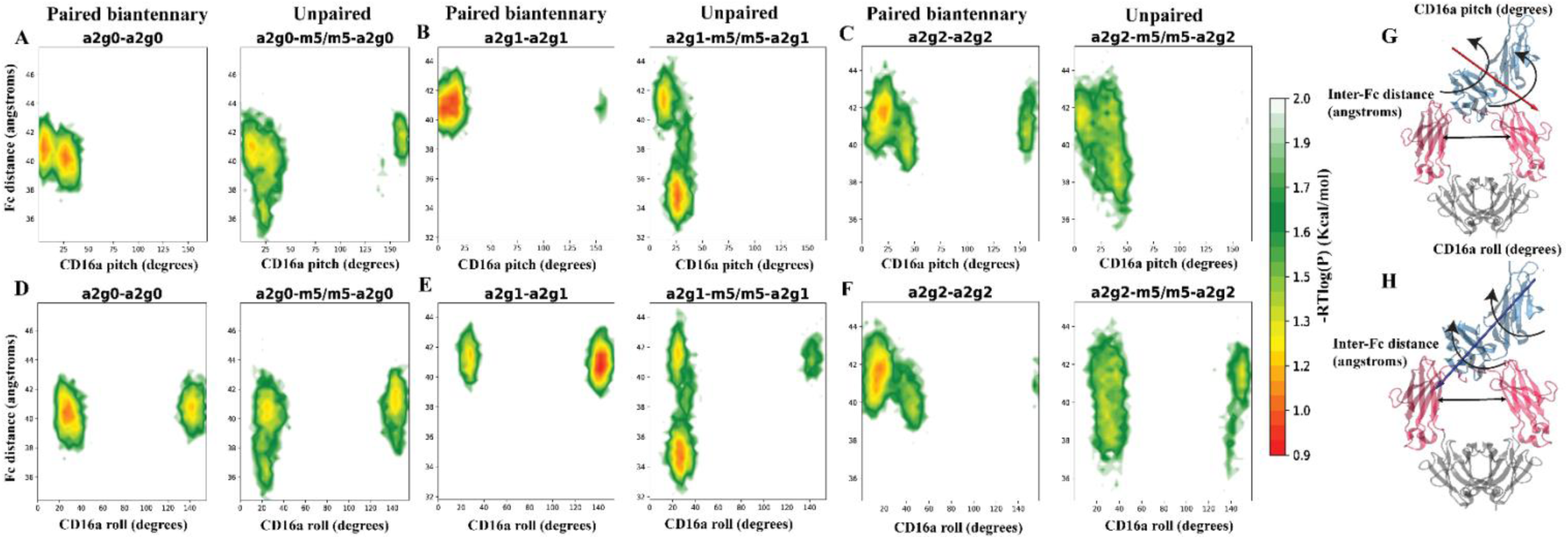
**Unpaired glycosylation destabilizes CD16a conformations relative to Paired biantennary glycosylation**. Free energy surface (FES) analysis of CD16a rotational motion along two independent axes; lower colorbar values indicate higher stability. **(A–F).** Each panel shows Paired biantennary glycosylation (left) and Unpaired glycosylation (right) **(G, H).** Rotational axes used for FES analysis. Paired biantennary systems sample compact conformational spaces with well-defined basins, while Unpaired systems exhibit diffuse ensembles with shallow basins.

Finally, we asked whether the dynamic and conformational differences between Paired Biantennary and Unpaired systems were accompanied by changes at the physical binding interface. Residue-wise energetic decomposition did not reveal major differences in the spatial distribution of energetic contributors between these two classes. We therefore quantified protein–protein contacts between Fc and CD16a using the g_contacts package of GROMACS^28, 29^. The contact maps showed that Paired Biantennary systems form substantially more protein–protein contacts than the corresponding Unpaired systems (**Figure 9A–C**), including approximately twice as many high-frequency contacts (with contact frequency greater than 0.6). Thus, although a single M5 glycan does not reorganize the residue-wise energetic landscape as strongly as Paired M5, it still measurably reduces the physical contacts that stabilize the Fc–CD16a interface.

**Figure 9:**
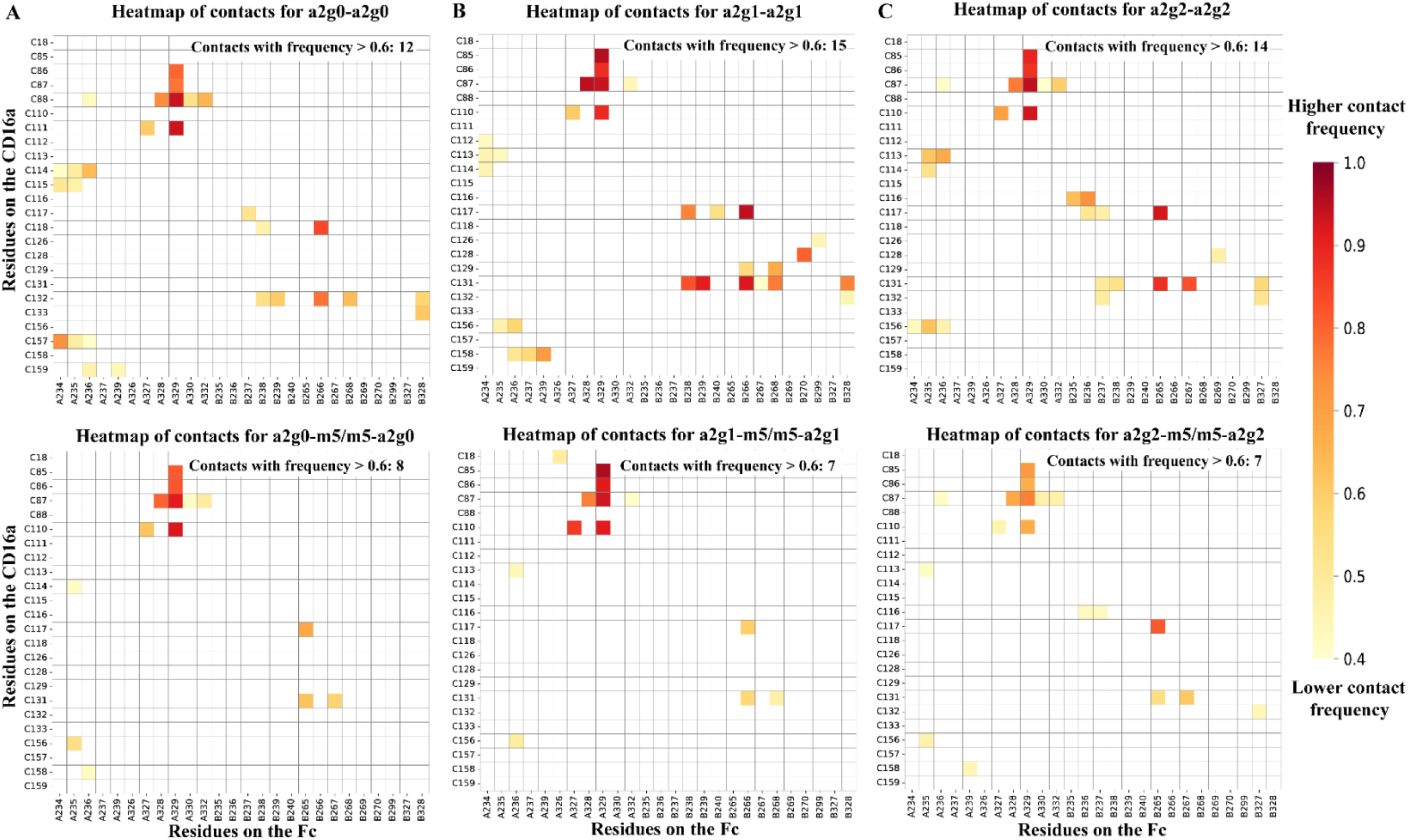
**Unpaired glycosylation reduces Fc–CD16a protein–protein contacts relative to Paired biantennary glycosylation**. Contact frequency analysis comparing Paired biantennary (top row) and Unpaired (bottom row) glycosylation on the Fc across **Panels A–C.** Paired biantennary systems show higher protein–protein contact frequencies than Unpaired variants, which could be leading to higher binding affinities.

**Figure 10:**
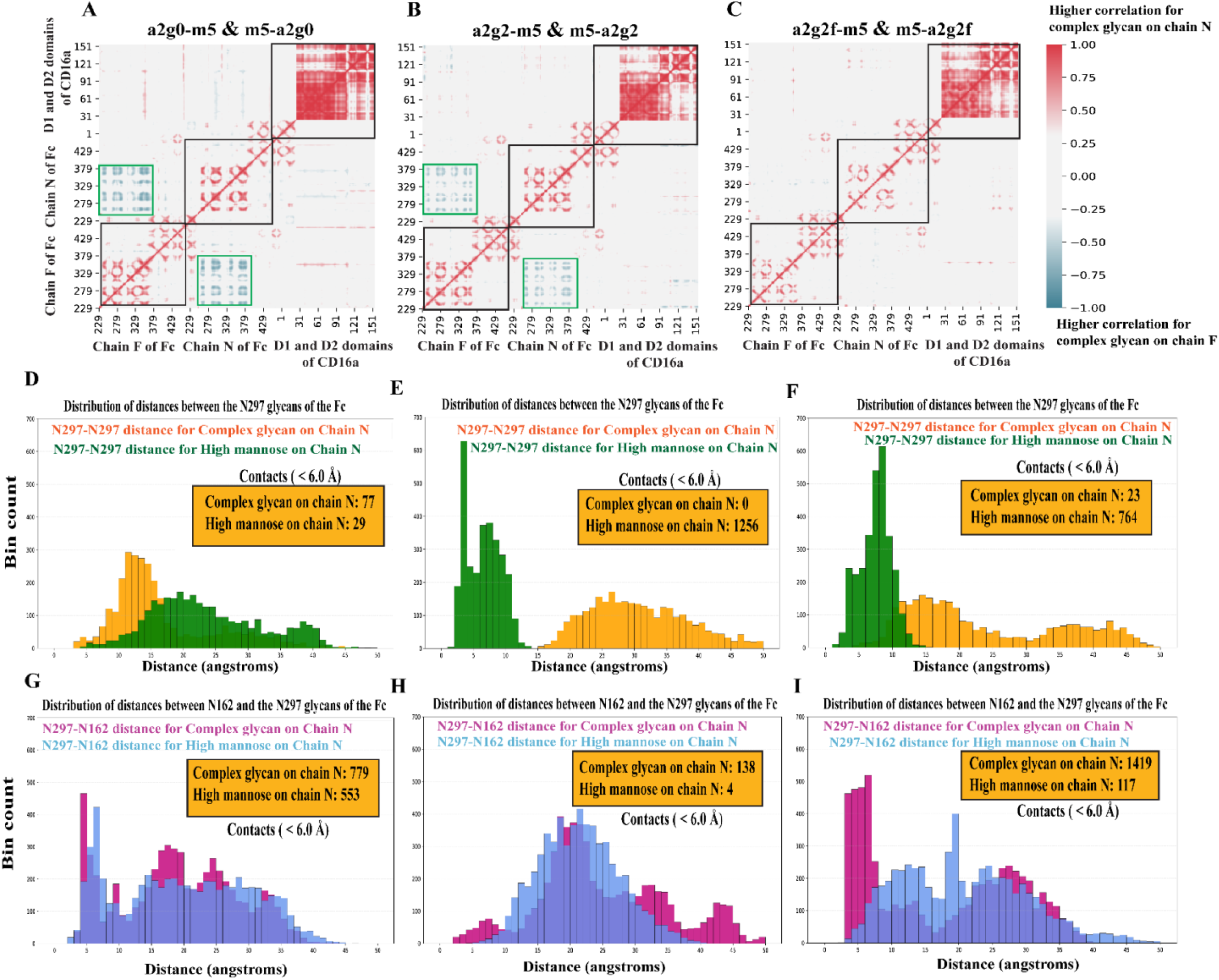
Proximal complex glycosylation enhances correlated domain motions, minimizes detrimental inter-glycan contacts, while maximizing productive inter-glycan contacts. (A, B, **C)** Dynamic cross-correlation analysis comparing the difference in correlated motions for Complex and High mannose glycosylation on chain N (the CD16a-proximal arm). Complex glycosylation on Chain N shows higher intra-Fc and intra-CD16a correlations (black squares), while high-mannose substitution shows increased inter-Fc correlations (green squares, **A** and **B**)**. (D, E, F)** Inter-glycan contact frequency analysis between the two Fc N297 glycans. The presence of a complex glycan on the proximal arm shows lower inter-Fc glycan contacts **(Panels E** and **F**), while higher contacts are seen in the case of **Panel D**, possibly due to the absence of galactose residues. (**G, H, I**) Distributions of distances between the Fc N297 glycan and the CD16a N162 glycan for complex glycan on chain N (magenta) versus high mannose on chain N (blue), for the same three systems. Complex glycan on chain N yields substantially more N297–N162 contacts than M5 in most systems, demonstrating that complex glycosylation on the CD16a-proximal Fc arm promotes tighter glycan–glycan packing at the Fc–CD16a interface, thereby aiding complex formation. This difference is substantial in **Panel I**, which could indicate its observed higher affinity of binding as higher productive contacts drive complex formation.

Together, these results show that a single M5 glycan is sufficient to impair productive Fc–CD16a recognition. Relative to Paired Biantennary glycosylation, Unpaired glycosylation reduces long-range dynamic coupling, broadens the receptor-bound conformational ensemble, and diminishes high-frequency protein–protein contacts at the interface. These findings suggest that the disruptive effect of M5 is not limited to symmetric Paired M5 glycosylation but can emerge from a single oligomannose substitution on one Fc arm. It should be noted, however, that these comparisons rely on energies averaged equally over both glycan-position combinations. In experimentally measured binding, the more energetically favorable positioning is likely to play a dominant role, so the equally weighted values may understate the contribution of glycan location to recognition.

### A complex glycan on the receptor-proximal Fc arm favors productive CD16a engagement

The preceding analyses showed that both Paired M5 and Unpaired glycosylation reduce Fc–CD16a binding relative to paired complex-type Fc glycosylation. However, because CD16a engages Fc asymmetrically, the functional effect of a single M5 glycan may depend not only on its presence, but also on which Fc arm it occupies. We therefore asked whether M5 on the receptor-proximal arm, Chain N, perturbs binding differently from M5 on the receptor-distal arm, Chain F. Experimentally, CD16a is known to bind the Fc asymmetrically^19, 30^. Azzam et.al. experimentally determined that in the case of mono-fucosylated glycans, the presence of the afucosylated glycan on the proximal arm leads to significantly higher binding affinities, which are comparable to those of doubly afucosylated systems^19^.

To test whether M5 positioning has an analogous effect, we compared theoretical binding energies for Unpaired systems in which the complex-type glycan or the M5 glycan occupied Chain N (**Figure 3B**). Across the systems analyzed, M5 on Chain N reduced binding affinity relative to the corresponding configuration in which the complex-type glycan occupied Chain N (**Figure 3B, Blocks 1–4**). The same trend was observed for the fucosylated comparison between a2g2f–m5 and m5–a2g2f (**Figure 3B, Block 4**), indicating that even in a higher-affinity fucosylated glycan context, receptor engagement is favored when the complex-type glycan is positioned on the CD16a-proximal Fc arm. Or as corollary, the receptor orients itself to engage Fc closer to the complex glycan arm.

We next examined the mechanistic basis for this positional dependence using correlated-motion and glycan-contact analyses (**Figure 10**). Systems with a complex-type glycan on Chain N showed stronger favorable correlated motions, particularly within the intra-Fc and intra-CD16a regions highlighted in **Figure 10A–C**. This suggests that a complex glycan on the receptor-proximal Fc arm promotes coordinated domain motions that support productive receptor engagement. In contrast, M5 on Chain N increased inter-Fc correlations in the green-boxed regions of **Figure 10A, 10B**, a pattern that is less consistent with productive Fc–CD16a coupling, as we saw in **Figure 7**. This unfavorable inter-Fc correlation pattern was absent in the fucosylated comparison in **Figure 10C**, providing a possible explanation for the comparatively higher binding affinity of the a2g2f–m5 system observed in **Figure 3B, Block 4**.

Inter-glycan contact analysis further supports this positional model. Increased contacts between the two Fc N297 glycans and the N162 glycan can drive favorable Fc–CD16a interactions. Consistent with this interpretation, systems with a complex-type glycan on the proximal Chain N minimized inter-Fc glycan contacts (**Figure 10E, 10F**), except in the case of a2g0-m5/m5-a2g0 which could be due to the absence of galactose residues (**Figure 10D**). However, increased contacts between the Fc glycan and the CD16a N162 glycan were observed in all cases (**Figure 10D–I**). In the case of an added fucose (**Panel F** and **I**), we observe the highest Fc-CD16a glycan contact frequency among the systems analyzed, indicating that the presence of a G2 fucosylated glycan on Chain N would lead to greater binding than that of a high-mannose.

Together, these results show that the effect of a single M5 glycan depends strongly on its position within the asymmetric Fc–CD16a interface. A complex-type glycan on the receptor-proximal Chain N promotes favorable correlated motions and Fc–CD16a glycan contacts, whereas M5 at the same position disrupts productive receptor engagement. These findings suggest that FcγRIIIa recognition is encoded not only by Fc glycan composition, but also by glycan placement, with the CD16a preferentially orienting towards the arm that has the complex glycan.

## DISCUSSION

Fc glycosylation is a central determinant of IgG1 effector function because it modulates the strength and geometry of Fcγ receptor engagement. While several Fc glycoengineering strategies, especially afucosylation, have been extensively studied and exploited to enhance FcγRIIIa/CD16a binding and ADCC activity^5, 31^, the structural consequences of high-mannose Fc glycans remain less clearly understood. This is particularly important because high-mannose glycans (most commonly M5) are frequently detected in therapeutic antibodies and are associated with accelerated serum clearance^32^. These observations highlight a central complexity: the effect of a glycan depends not only on its chemical identity, but also on whether it is located on the antibody and where it sits within the asymmetric Fc–CD16a interface. Here, we used all-atom MD simulations to define how oligomannosidic M5 glycans on IgG1 Fc alter CD16a recognition. Our simulations reproduced the experimentally observed energetic hierarchy among Fc glycoforms: Paired M5 systems showed weaker Fc–CD16a interaction energies than Paired Biantennary or Unpaired systems, consistent with experimental measurements showing reduced CD16a binding for M5-containing Fc glycoforms relative to complex-type paired glycans^19, 20, 30^. This agreement provides confidence that the simulations capture the relevant glycoform-dependent binding profiles and supports using the trajectories to interrogate the structural and dynamic origins of these differences. Importantly, this reduction was not simply a uniform energetic weakening of the interface. Residue-wise decomposition showed that Paired M5 shifts energetically active CD16a residues away from the productive Fc-contacting surface, while contact analyses showed reduced protein–protein and glycan-mediated physical contacts across the receptor-bound complex. Thus, M5 glycosylation changes both the energetic and structural organization of Fc–CD16a recognition. Second, our conformational and dynamic analyses show that the effect of M5 extends beyond the local binding interface. Free energy surface analyses revealed that Paired Biantennary and Unpaired systems occupy more compact, stable receptor-bound basins, whereas Paired M5 systems sample broader and more heterogeneous conformational landscapes. Structures from these basins suggest that productive CD16a engagement is associated with receptor orientations that maintain favorable proximity between the D2 domain and the receptor-proximal Fc arm. In contrast, Paired M5 shifts the complex toward less well-defined binding geometries. Dynamic cross-correlation analysis further showed that Paired M5 suppresses intra-Fc, intra-CD16a, and inter-domain Fc–CD16a coupling. Although M5-containing systems can exhibit enhanced correlations between the two Fc arms, these motions appear unproductive because they are not accompanied by stronger Fc–CD16a coupling or improved receptor engagement. Notably, even the presence of a single M5 glycan increases variation in Fc-arm separation, indicating that asymmetric oligomannose substitution is sufficient to perturb the relative positioning of the two Fc arms. These findings suggest that productive receptor engagement requires not only favorable contacts but also coordinated motions across the Fc–CD16a complex.

A third major finding is that a single M5 glycan is sufficient to perturb receptor recognition. Previous work showed that Paired Biantennary G0 glycoforms bind more strongly than corresponding Unpaired G0–M5 systems, although the glycoforms studied were much more limited^19^. Our simulations show that Paired Biantennary systems consistently exhibit stronger computed binding than Unpaired counterparts across the glycoforms examined. Mechanistically, Unpaired systems showed reduced long-range dynamic coupling, broader conformational sampling, and fewer high-frequency protein–protein contacts compared with Paired Biantennary systems. Therefore, the disruptive effect of M5 is not restricted to symmetric high-mannose glycosylation on both Fc arms; even a single oligomannose substitution can alter the physical and dynamic organization of the receptor-bound complex.

The final insight from this study is that glycan position matters. CD16a binds Fc asymmetrically, and prior studies have shown that asymmetric Fc engineering or glycosylation can markedly alter receptor binding and ADCC activity^16^. Azzam et al. further showed that an afucosylated glycan on the receptor-proximal Chain N can produce affinities comparable to fully afucosylated systems^19^. Our results reveal a parallel positional principle for M5: a complex-type glycan on Chain N favors productive CD16a engagement, whereas M5 on Chain N is particularly disruptive. Systems with complex glycans on the proximal arm showed more favorable correlated motions and increased Fc–CD16a glycan contacts involving the CD16a N162 glycan, while minimizing inter-Fc glycan contacts that could sterically constrain the interface. These results suggest the Fc arm carrying the glycan composition most compatible with productive receptor engagement is a strong determinant of preferential orientation of CD16a. However, this orientational preference is not expected to be determined by glycan identity alone. It is likely shaped by other sources of asymmetry as well, including the initial receptor-Fc encounter pose, hinge and CH2-domain fluctuations, local glycan conformations, and transient contacts with CD16a glycans such as N162.

Some limitations should be considered when interpreting these results. Although the simulations were performed in three independent microsecond-scale replicates for each system, they were initiated from the only known Fc–CD16a structural orientation and therefore may not capture the full diversity of receptor-binding geometries accessible experimentally. One must also keep in mind that CD16a glycosylation was represented using the most probable receptor glycoforms, whereas CD16a glycosylation is heterogeneous and can itself modulate Fc binding to some degree. In addition, the energies reported for the Unpaired systems reflect an equal (50:50) weighting of the two glycan-position combinations rather than their true relative populations. This is because energetically unfavorable combinations are likely to be less frequented and short-lived, and the more favorable positioning is expected to dominate actual binding events. While, this equal weighting may not be fully representative of the experimentally measured energetics, further exploration of the Free Energy Surface (FES) results along with experimental binding energies, can be used to recover position-specific binding energies and to weigh the two combinations by their true relative ratios. Also, extending this framework to additional Fc and CD16a glycoforms, Fcγ receptor variants, and engineered Fc mutants would further clarify how glycan composition and protein-interface residues jointly tune effector function.

The findings in this study have important practical implications for antibody design. Glycoengineering strategies should consider not only the average glycan composition of an antibody preparation, but also the pairing and arm-specific placement of glycans within the Fc dimer. Choice of symmetric/asymmetric Fc glycosylation, combined with residue-level information from energetic decomposition and contact-map analyses, could be used to tune the probability of productive receptor-binding orientations. Such an approach would allow glycan composition and protein-interface engineering to be considered together, with the goal of stabilizing desired Fc–CD16a engagement modes while avoiding glycoforms associated with rapid clearance. More broadly, this work illustrates how simulations can complement glycoprofiling and biochemical assays by resolving how specific glycan arrangements encode receptor recognition, providing structural design rules for therapeutic antibodies with tunable effector function.

## CONCLUSION

This study provides a molecular framework for understanding how oligomannosidic M5 glycosylation on IgG1 Fc modulates CD16a recognition. Across a panel of Paired Biantennary, Unpaired, and Paired M5 glycoforms, our simulations show that M5-containing Fc glycans reduce productive Fc–CD16a engagement in a manner that depends on both glycan pairing and receptor-bound orientation. Paired M5 glycosylation produces the strongest disruption, weakening binding energetics, redistributing energetic contributions away from the productive interface, reducing protein–protein and glycan-mediated physical contacts, and destabilizing receptor-bound conformational ensembles. Notably, a single M5 glycan is also sufficient to perturb the complex, increasing Fc-arm separation heterogeneity and reducing the correlated motions and high-frequency contacts associated with stronger receptor engagement. Together, these findings suggest that FcγRIIIa recognition is encoded not only by Fc glycan composition, but also by the combined effects of glycan pairing, conformational orientation, and protein-interface contacts. This has direct implications for antibody design: asymmetric Fc glycosylation and residue-level engineering, guided by energetic decomposition and contact-map analyses, may be used together to bias Fc–CD16a complexes toward productive binding modes. More broadly, this work illustrates how molecular simulations can resolve glycan-dependent mechanisms that are difficult to isolate experimentally, providing design principles for tuning therapeutic antibody effector function.

## METHODS

The crystal structure of the IgG1 Fc-CD16a complex (code: 1E4K) was obtained from RCSB^20^. Residue numbering throughout this study follows that of the crystal structure (PDB ID: 1E4K), with the Fc domain spanning residues 229–444 in each chain. The Fc domain of the complex was glycosylated with different degrees of galactosylation (G0, G1 and G2) in this study, for both Paired Biantennary and Unpaired systems (see **Panel B** of **Figure 2** and **SI Table 1**). CD16a glycans were modelled from the most probable glycoforms in the mass-spectrometry data of Kashyap et al., with the N162 glycan replaced by high-mannose (M5) to elicit the strongest binding^5, 9^. All systems are listed in **SI Table 1**. Glycans were added to the crystal structure using ALLOSMOD, an ab initio modelling tool^33^, with the CHARMM36 force field^34^. For Unpaired systems, the notation a2gx-m5 (x = 0, 1, 2) indicates that the complex glycan occupies the proximal arm and the mannose the distal arm; the reverse notation applies for the opposite arrangement. All the results for these Unpaired systems are from the combined simulations of each of the pairs and the energies reported for the Unpaired systems were obtained as a simple (50:50) average of the two complementary glycan-position combinations (i.e., a2gx-m5 and m5-a2gx). The final structure comprises Chain N and Chain F of the IgG1 Fc with 13 distinct glycoforms, bound to the D1 and D2 domains of CD16a, which carries the highest-occupancy glycans modelled at N38, N45, N74, and N169. Visualisations and structural representations were prepared with VMD 1.9.335 and PyMOL^36^

### Simulation protocol

We had a total of 13 distinct systems as represented in **SI Table 1**. A transferable intermolecular potential with 3 points (TIP3P) water model^37^ was used for simulations, and a cubic waterbox was used as the simulation box with a padding of 20 Å to account for the motion and possible extension of glycans. The systems were neutralised at a physiological ionic concentration of 150 mM of KCl. Since we used CHARMM36 forcefield, for best forcefield compatibility and structural setup, NAMD was used for creating topologies^38^. Energy minimisation was performed in NAMD using combined conjugate-gradient and steepest-descent algorithms, with constraints on glycan heavy atoms and the protein backbone gradually released. To harness the optimum graphical processing unit (GPU) enabled speedup, the topology and parameter files for these minimized systems were converted to AMBER form^25^, where all the equilibration production simulations were performed. This also allowed us to use the hydrogen mass repartitioning capability in AMBER, which allows a simulation timestep of 4 fs, and helps to speed up the computation time^39^.

The first equilibration step involved a restrained simulation in a constant volume and temperature (NVT) ensemble for 2 ns. The second step involved a restrained simulation with constant pressure and temperature (NPT) for 10 ns. In both stages, harmonic position restraints were applied to all non-hydrogen atoms. The third step of equilibration involved 10 ns of simulation in the NPT ensemble and only included position restraints on protein backbone atoms.

After this equilibration protocol, we ran three separate trajectories for each system, each around 1.2 μs in length. Temperature was maintained at 310 K using a Langevin thermostat (canonical ensemble sampling). Pressure was maintained at 1 atm using a Berendsen barostat. This combination was used for all production trajectories. Upon examining the RMSD plots, the first 200 ns of each simulation were discarded to account for the equilibration of the system (see **SI Figure 1**). Hence, the total simulation time was 3 μs for each system.

The Paired M5 system required an additional enhanced-sampling step before atomistic MD, because the IgG1 Fc–CD16a crystal structure (PDB 1E4K)^20^ carries complex glycans on both arms and therefore does not capture the bound conformation under M5 glycosylation. The minimized Paired-M5 complex was subjected to accelerated MD (aMD) using boost parameters computed per Pierce et al.^40^. Dihedral and local torsions were boosted to prevent large-scale protein unfolding. After 100 ns of aMD, the final structure was used as the starting point for atomistic MD as described above.

### Estimation of theoretical binding energies

Binding energy calculations were performed using the MMPBSA.py package^24^ of AMBER^25^. Frames from the final 600 ns of each trajectory were used (1800 frames total across the three trajectories). We calibrated the relative binding energies using an empirically corrected MM-PBSA approach, in which the van der Waals and electrostatic contributions are retained but scaled by literature-derived coefficients (α = 0.158 for vdw and β = 0.153 for electrostatic, respectively) to reduce the well-known tendency of single-trajectory continuum-solvent methods to overestimate absolute interaction energies, especially for buried charged contacts and protein–protein interfaces, following the approach of Li et al.^26^. This preserves the efficiency and comparative interpretability of MM-PBSA while placing affinities on a more experimentally realistic scale, making the values suitable for comparison across glycosylation profiles rather than as absolute free-energy estimates. The use of small scaling factors for these terms is also conceptually consistent with earlier linear interaction energy (LIE)-like formulations, where raw interaction energies are empirically damped to account for solvent reorganisation and structural relaxation not explicitly captured in simplified free-energy models^41^. In the case of Unpaired systems, where different glycans were placed on the two homologous chains of Fc, we calculated the average values of the Van der Waals and Electrostatic energy components, which were taken from these systems, along with their standard errors.

### Analysis of glycan-glycan interactions and protein-protein contacts

Glycan–glycan interactions were quantified for two cases: (i) between the heavy atoms of the two N297 Fc glycans, and (ii) between the heavy atoms of the two N297 Fc glycans and the N162 CD16a glycan. The minimum heavy atom distance in both cases was computed per frame using all pairwise heavy atom combinations, and the average minimum distance across the trajectory was used to assess differences in glycan–glycan proximity. For case (ii), the minimum distances from each of the two N297 Fc glycans to the N162 glycan were computed independently and combined across both Fc arms to give a composite measure of N297–N162 proximity. Contact frequency values within 6.0 Å were considered favourable contacts. This cutoff is consistent with standard heavy-atom contact definitions used in structural analyses^42^ and prior Fc glycan simulation studies^43^.

Protein–protein contacts between Fc and CD16a were computed with the g_contacts module of GROMACS using a default cutoff of 3.0 Å^28, 29^. Contact frequencies below 0.25 were discarded, and inter-system differences were plotted instead of absolute frequencies to highlight the changes between systems.

### Residue-wise energy decomposition analysis

Per-residue energy decomposition was used to identify residues contributing most to the Fc–CD16a interface, performed with the MMPBSA.py package^24^ of AMBER^25^. A portion of the trajectories was taken as the input, specifically the frames corresponding to the time between 600 and 1200 ns, yielding a total of 50 evenly spaced frames for each system. The 20 residues with the largest contributions to the binding free energy at the interface were identified for each system. To compare Paired Biantennary/Unpaired and Paired-M5 complexes, residues uniquely enriched in each were mapped onto the Fc–CD16a structures, allowing the spatial distribution of energetically critical residues to be visualized and quantified across the interface.

### Analysis of the motion of the CD16a relative to the Fc

To characterize the impact of glycans on the motion of CD16a relative to the Fc domain, we quantified both rotational and translational degrees of freedom of these two domains. This was calculated by measuring the relative orientation of CD16a and was described using two principal axes of rotation (see **Panels E** and **F** of **Figure 6, Panels G** and **H** of **Figure 8 and Panels I and J** of **SI Figure 4**). In case of the Fc, two distinct motions were identified: (i) separation within the two arms, which were monitored by measuring the distance between the Centre of mass (COM) of the two CH2 domains and (ii) opening/closing of the two arms relative to the CD16a domain, which was calculated by measuring the angle formed between the COM of each of CH1 and CH2 domains of the Fc with respect to the CD16a domain. These reaction coordinates were chosen to capture coupled motions between CD16a reorientation and Fc conformational changes.

The conformational landscapes associated with these motions were quantified using two-dimensional Free Energy Surface (FES) plots. Probability distributions along each pair of reaction coordinates were extracted from equilibrated simulation trajectories and converted to free-energy surfaces. In these FES representations, deeper energy basins correspond to more stable and frequently sampled conformational states.

### Analysis of Fc-CD16a correlated motions

Dynamic cross-correlation (DCC) analysis was used to probe the correlative motions of atoms in the Fc and the CD16a. This correlative motion between two atoms *i* and *j* is defined as

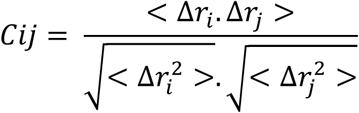

where *ri*(*t*) denotes the vector of the *i*th atom’s coordinates as a function of time *t*, < >t is the time ensemble average and Δ*ri*(t) = *ri*(*t*) − < *ri*(*t*)*T*>*t*. The difference in the values for the afucosylated and fucosylated systems is then plotted in the form of an *N* × *N* heatmap, where *N* is the number of Cα atoms in the system. The correlation values are calculated between −1 and 1, where 1 = complete correlation; −1 = complete anti-correlation; 0 = no correlation. The DCC matrix was generated using the MD-Task^27^ Python library and plotted using MatPlotLib^44^.

All data analyses and simulation interpretations were performed using the Python programming language, including the MDtraj library for trajectory analyses^45^, MD-TASK for dynamic cross-correlation analysis^27^ and Matplotlib^44^ for visualization.

### Supplementary material

The Supplementary material contains figures that are relevant to the findings in the Main manuscript. These include the complete list of systems, RMSD plots, Energy decomposition plots. protein-protein and glycan-glycan contact analysis, Free energy surface (FES) analysis and Dynamic cross correlation (DCC) plots.

Filename: “**Supplementary Information**”

## Funding and Acknowledgements

N.M. and S.C. were supported by NIH NIGMS grant R35GM151231-01 and through NEU Faculty Startup Funds. N. M. acknowledges the LEADERS fellowship program jointly sponsored by Northeastern University and Amgen. This research used computational resources from the Northeastern Discovery cluster at MGHPCC. The authors also acknowledge the initial tests performed by Raghavendran Suresh, which led to the start of this project.

## Author contributions

N.M. A.P. and S.C. conceptualized the study and designed the experiments. N.M. and S.C. prepared the initial structures. N.M. ran the simulations. N.M. performed the data analysis. N.M., A.P. and S.C. performed data interpretation. N.M. prepared the figures. N.M, A.P. and S.C wrote and edited the manuscript. S.C. is the corresponding author.

## Competing interests

The authors declare no competing interests.

## Data and materials availability

Simulation trajectories and data are shared for open access in doi: 10.17632/6ncyp3j9sy.1. Further materials and simulation codes are available from the corresponding authors upon request.

## Notes

### Competing Interest Statement

The authors have declared no competing interest.

## REFERENCES

1. Vidarsson, G., Dekkers, G. & Rispens, T. IgG subclasses and allotypes: from structure to effector functions. Frontiers in Immunology 5, 520 (2014).

2. Gessner, J.E., Heiken, H., Tamm, A. & Schmidt, R.E. The IgG Fc receptor family. Ann Hematol 76, 231–248 (1998).

3. Robert L. Shields, J.L., Rodney Keck, Lori Y. O’Connell, Kyu Hong, Y. Gloria Meng, & Stefanie H. A. Weikert, a.L.G.P. Lack of Fucose on Human IgG1 N-Linked Oligosaccharide Improves Binding to Human Fc RIII and Antibody-dependent Cellular Toxicity. Journal of Biological Chemistry (2002).

4. Falconer, D.J., Subedi, G.P., Marcella, A.M. & Barb, A.W. Antibody Fucosylation Lowers the FcgammaRIIIa/CD16a Affinity by Limiting the Conformations Sampled by the N162-Glycan. ACS Chem Biol 13, 2179–2189 (2018).

5. Subedi, G.P. & Barb, A.W. CD16a with oligomannose-type N-glycans is the only “low-affinity” Fc gamma receptor that binds the IgG crystallizable fragment with high affinity in vitro. Journal of Biological Chemistry 293, 16842–16850 (2018).

6. Rodriguez Benavente, M.C., Hughes, H.B., Kremer, P.G., Subedi, G.P. & Barb, A.W. Inhibiting N-glycan processing increases the antibody binding affinity and effector function of human natural killer cells. Immunology 170, 202–213 (2023).

7. Barb, G.P.S.a.A.W. The immunoglobulin G1 N-glycan composition affects binding to each low affinity Fcg receptor. MABS (2016).

8. Jacob T. Roberts, K.R.P., and Adam W. Barb Site-specific N-glycan Analysis of Antibody-binding Fcg Receptors from Primary Human Monocytes. Molecular & Cellular Proteomics (2020).

9. Benavente, M.C.R., Patel, K.R., Lorenz, W.W., Mace, E.M. & Barb, A.W. Fc γ receptor IIIa/CD16a processing correlates with the expression of glycan-related genes in human natural killer cells. Glycobiology 30, 1102–1102 (2020).

10. Delidakis, G., Kim, J.E., George, K. & Georgiou, G. Improving Antibody Therapeutics by Manipulating the Fc Domain: Immunological and Structural Considerations. Annu Rev Biomed Eng 24, 249–274 (2022).

11. Julie Van Coillie, M.A.S., Arthur E. H. Bentlage, Noortje de Haan, Zilu Ye, Dionne M. Geerdes, Wim J. E. van Esch, Lise Hafkenscheid, Rebecca L. Miller, Yoshiki Narimatsu, Sergey Y. Vakhrushev, Zhang Yang, Gestur Vidarsson and Henrik Clausen Role of N-Glycosylation in FcgRIIIa interaction with IgG. frontiers in Immunology (2022).

12. Hajduk, J. et al. Interaction analysis of glycoengineered antibodies with CD16a: a native mass spectrometry approach. MAbs 12, 1736975 (2020).

13. Kiyoshi, M. et al. Assessing the Heterogeneity of the Fc-Glycan of a Therapeutic Antibody Using an engineered FcgammaReceptor IIIa-Immobilized Column. Scientific Reports 8, 3955 (2018).

14. Liu, Y.D. & Flynn, G.C. Effect of high mannose glycan pairing on IgG antibody clearance. Biologicals 44, 163–169 (2016).

15. Goetze, A.M. et al. High-mannose glycans on the Fc region of therapeutic IgG antibodies increase serum clearance in humans. Glycobiology 21, 949–959 (2011).

16. Meudt, M. et al. Spotlight on Glycan Pairing: The Generation and Impact of Monoclonal Antibody Asymmetrical Fc N–Glycan Pairs on Fc Receptor Interaction. ACS Pharmacol Transl Sci 8, 1756–1767 (2025).

17. Hayes, J.M. et al. Fc Gamma Receptor Glycosylation Modulates the Binding of IgG Glycoforms: A Requirement for Stable Antibody Interactions. Journal of Proteome Research 13, 5471–5485 (2014).

18. Lampros, E.A. et al. The antibody-binding Fc gamma receptor IIIa / CD16a is N-glycosylated with high occupancy at all five sites. Current Research in Immunology 3, 128–135 (2022).

19. Azzam, T. et al. Asymmetrically glycosylated IgG1 antibodies are universal and drive human disease. Nature Communications 17, 383 (2025).

20. Peter Sondermann, R.H., Vaughan Oosthuizen & Uwe Jacob The 3.2-A crystal structure of the human IgG1 Fc fragment±FcgRIII complex. nature (2000).

21. X Li, T.P., EE Brown and JC Edberg Fcg receptors: structure, function and role as genetic risk factors in SLE. Genes and Immunity (2009).

22. Drescher, B., Witte, T. & Schmidt, R.E. Glycosylation of FcgammaRIII in N163 as mechanism of regulating receptor affinity. Immunology 110, 335–340 (2003).

23. Huang, J. et al. CHARMM36m: an improved force field for folded and intrinsically disordered proteins. Nat Methods 14, 71–73 (2017).

24. Miller, B.R., 3rd et al. MMPBSA.py: An Efficient Program for End-State Free Energy Calculations. Journal of Chemical Theory and Computation 8, 3314–3321 (2012).

25. Case, D.A. et al. AmberTools. Journal of Chemical Information and Modeling 63, 6183–6191 (2023).

26. Li, M. & Zheng, W. Probing the structural and energetic basis of kinesin-microtubule binding using computational alanine-scanning mutagenesis. Biochemistry 50, 8645–8655 (2011).

27. Brown, D.K. et al. MD-TASK: a software suite for analyzing molecular dynamics trajectories. Bioinformatics 33, 2768–2771 (2017).

28. Blau, C. & Grubmuller, H. g_contacts: Fast contact search in bio-molecular ensemble data. Computer Physics Communications 184, 2856–2859 (2013).

29. Bekker, H. et al. Gromacs - a Parallel Computer for Molecular-Dynamics Simulations. Physics Computing *’*92 **92**, 252–256 (1993).

30. Liu, Z. et al. Asymmetrical Fc engineering greatly enhances antibody-dependent cellular cytotoxicity (ADCC) effector function and stability of the modified antibodies. Journal of Biological Chemistry 289, 3571–3590 (2014).

31. Kremer, P.G., Lampros, E.A., Blocker, A.M. & Barb, A.W. One N-glycan regulates natural killer cell antibody-dependent cell-mediated cytotoxicity and modulates Fc gamma receptor IIIa/CD16a structure. Elife 13, RP100083 (2024).

32. Pereira, N.A., Chan, K.F., Lin, P.C. & Song, Z. The “less-is-more” in therapeutic antibodies: Afucosylated anti-cancer antibodies with enhanced antibody-dependent cellular cytotoxicity. MAbs 10, 693–711 (2018).

33. Guttman, M., Weinkam, P., Sali, A. & Lee, K.K. All-atom ensemble modeling to analyze small-angle x-ray scattering of glycosylated proteins. Structure 21, 321–331 (2013).

34. Brooks, B.R. et al. CHARMM: the biomolecular simulation program. J Comput Chem 30, 1545–1614 (2009).

35. Humphrey, W., Dalke, A. & Schulten, K. VMD: visual molecular dynamics. J Mol Graph 14, 33–38, 27-38 (1996).

36. 36. Schrödinger, L., & DeLano, W. PyMOL. (2020).

37. Jorgensen, W.L., Chandrasekhar, J., Madura, J.D., Impey, R.W. & Klein, M.L. Comparison of Simple Potential Functions for Simulating Liquid Water. Journal of Chemical Physics 79, 926–935 (1983).

38. Phillips, J.C. et al. Scalable molecular dynamics on CPU and GPU architectures with NAMD. The Journal of Chemical Physics 153, 044130 (2020).

39. Hopkins, C.W., Le Grand, S., Walker, R.C. & Roitberg, A.E. Long-Time-Step Molecular Dynamics through Hydrogen Mass Repartitioning. Journal of Chemical Theory and Computation 11, 1864–1874 (2015).

40. Pierce, L.C., Salomon-Ferrer, R., Augusto, F.d.O.C., McCammon, J.A. & Walker, R.C. Routine Access to Millisecond Time Scale Events with Accelerated Molecular Dynamics. Journal of Chemical Theory and Computation 8, 2997–3002 (2012).

41. Carlsson, J., Ander, M., Nervall, M. & Aqvist, J. Continuum solvation models in the linear interaction energy method. The Journal of Physical Chemistry B 110, 12034–12041 (2006).

42. Yuan, C., Chen, H. & Kihara, D. Effective inter-residue contact definitions for accurate protein fold recognition. BMC Bioinformatics 13, 292 (2012).

43. Lee, H.S. & Im, W. Effects of N-Glycan Composition on Structure and Dynamics of IgG1 Fc and Their Implications for Antibody Engineering. Scientific Reports 7, 12659 (2017).

44. Hunter, J.D. Matplotlib: A 2D Graphics Environment. Computing in Science & Engineering 9, 90–95 (2007).

45. McGibbon, R.T. et al. MDTraj: A Modern Open Library for the Analysis of Molecular Dynamics Trajectories. Biophys J 109, 1528–1532 (2015).

