## Supplemental Information Tables and Figures for "Oligomannose Fc Glycans Reprogram the Energetic and Conformational Basis of CD16a Recognition"

| System ID | Glycan on chain N | Glycan on chain F | Type |
|---|---|---|---|
| a2g0-a2g0 | a2g0 | a2g0 | Paired biantennary |
| a2g1-a2g1 | a2g1 | a2g1 | Paired biantennary |
| a2g2-a2g2 | a2g2 | a2g2 | Paired biantennary |
| a2g2f-a2g2f | a2g2f | a2g2f | Paired biantennary |
| a2g0-m5 | a2g0 | m5 | Unpaired |
| m5-a2g0 | m5 | a2g0 | Unpaired |
| a2g1-m5 | a2g1 | m5 | Unpaired |
| m5-a2g1 | m5 | a2g1 | Unpaired |
| a2g2-m5 | a2g2 | m5 | Unpaired |
| m5-a2g2 | m5 | a2g2 | Unpaired |
| a2g2f-m5 | a2g2f | m5 | Unpaired |
| m5-a2g2f | m5 | a2g2f | Unpaired |
| m5-m5 | m5 | m5 | Paired high mannose |

**SI Table 1:** The table shows the 13 different glycan structural variants analyzed in our study and categorized by glycan composition (Paired biantennary, Unpaired and Paired High mannose). Chain N and Chain F are the CD16a proximal and distal arms of the Fc, respectively.

| System | Residue on the Fc | Location on the Fc | Residue on the CD16a | Location on the CD16a |
|---|---|---|---|---|
| a2g1-m5/m5-a2g1 | PRO329 | Chain F | TRP110 | D2 |
| a2g1-m5/m5-a2g1 | LEU223 | Chain N | ALA114 | D2 |
| a2g1-m5/m5-a2g1 | LYS288 | Chain F | LEU154 | D2 |
| a2g1-m5/m5-a2g1 | LYS326 | Chain F | TRP87 | D2 |
| a2g1-m5/m5-a2g1 | LEU222 | Chain N | HSE116 | D2 |
| a2g1-m5/m5-a2g1 | TRP277 | Chain F | LYS158 | D2 |
| a2g1-m5/m5-a2g1 | ASN276 | Chain F | HSE131 | D2 |
| a2g1-m5/m5-a2g1 | LEU328 | Chain F | HSE142 | D2 |
| a2g1-m5/m5-a2g1 | LYS274 | Chain F | ILE85 | D1 |
| a2g1-m5/m5-a2g1 | ASP258 | Chain N | LYS117 | D2 |
| a2g1-a2g1 | PRO329 | Chain F | GLU82 | D1 |
| a2g1-a2g1 | ASP253 | Chain N | TRP110 | D2 |
| a2g1-a2g1 | LYS326 | Chain F | TYR129 | D2 |
| a2g1-a2g1 | PRO220 | Chain N | VAL118 | D2 |
| a2g1-a2g1 | LEU235 | Chain F | GLU163 | D2 |
| a2g1-a2g1 | LEU234 | Chain F | LEU115 | D2 |
| a2g1-a2g1 | ALA327 | Chain F | ALA114 | D2 |
| a2g1-a2g1 | LEU328 | Chain F | LEU154 | D2 |
| a2g1-a2g1 | PRO238 | Chain F | PHE130 | D2 |
| a2g1-a2g1 | ASP265 | Chain F | TRP87 | D2 |
| m5-m5 | ASP253 | Chain N | LYS19 | D1 |
| m5-m5 | ASP270 | Chain F | ARG152 | D2 |
| m5-m5 | ASP269 | Chain F | LYS4 | D1 |
| m5-m5 | ASP293 | Chain F | GLY156 | D2 |

| | | | | |
|---|---|---|---|---|
| m5-m5 | SER227 | Chain N | LYS140 | D2 |
| m5-m5 | ASP280 | Chain F | ARG67 | D1 |
| m5-m5 | GLU333 | Chain F | LYS144 | D2 |
| m5-m5 | ASP265 | Chain F | SER157 | D2 |
| m5-m5 | GLU318 | Chain F | LYS111 | D2 |
| m5-m5 | GLU258 | Chain N | LYS98 | D2 |

**SI Table 2** The top ten energetically active residues identified at the Fc–CD16a interface for each glycoform complex, categorized by Fc chains and CD16a domains. The entire list can be provided upon request. Note that the residue numbering throughout this study follows that of the crystal structure (PDB ID: 1E4K), with the Fc domain spanning residues 229–444 in each chain.

### RMSD plots

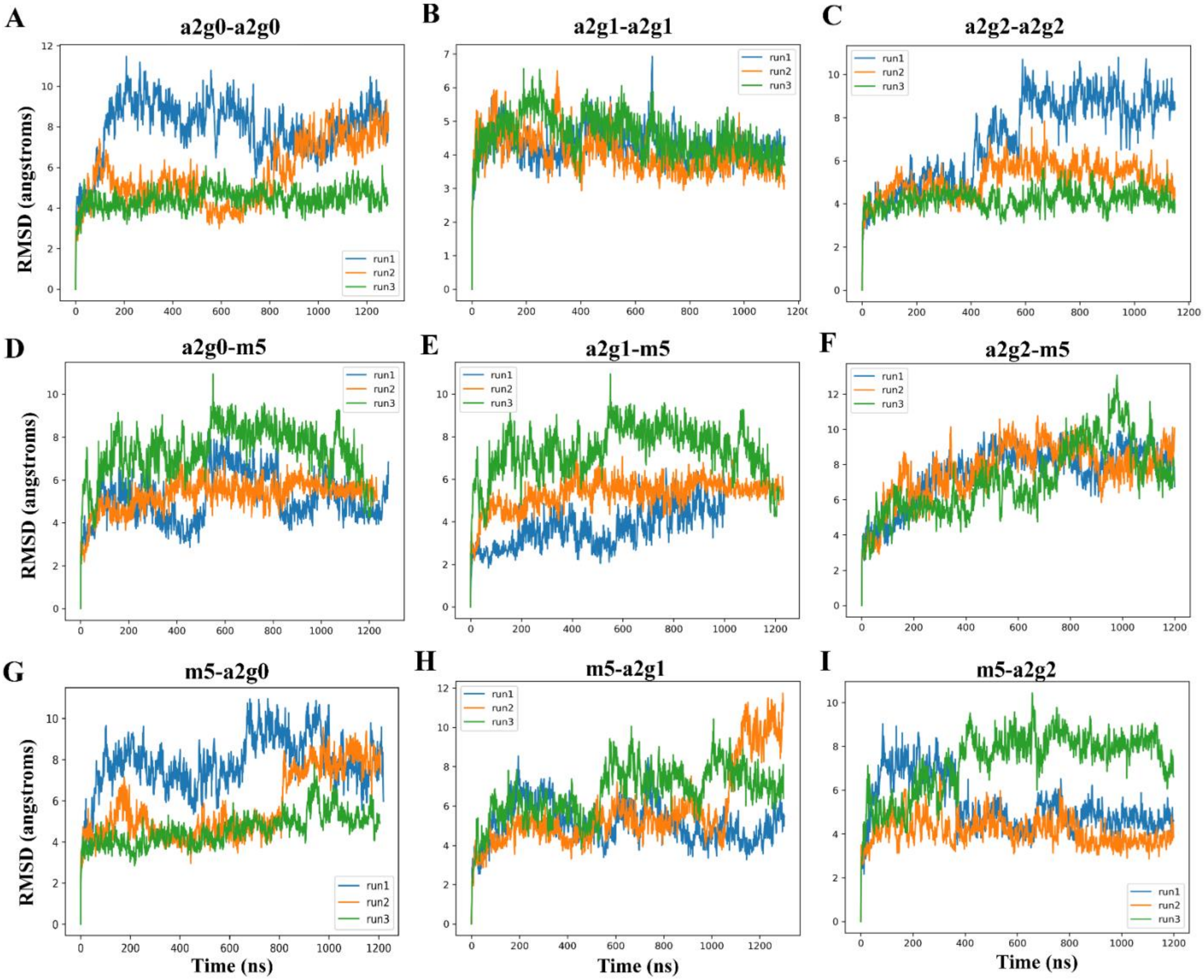


**SI Figure 1**: RMSD analysis demonstrates that equilibrium around local minima is reached beyond 200ns and generally remain there for each trajectory across select glycosylated Fc-CD16a systems. The RMSD (in angstroms) of the Cα atoms is plotted as a function of simulation time (in ns) for three independent replicates (run1, run2 and run3) of each Paired biantennary (**Panels A**, **B** and **C**), Unpaired with M5 on chain F (**Panels D**, **E** and **F**) and Unpaired with m5 on chain N (**Panels G**, **H** and **I**).

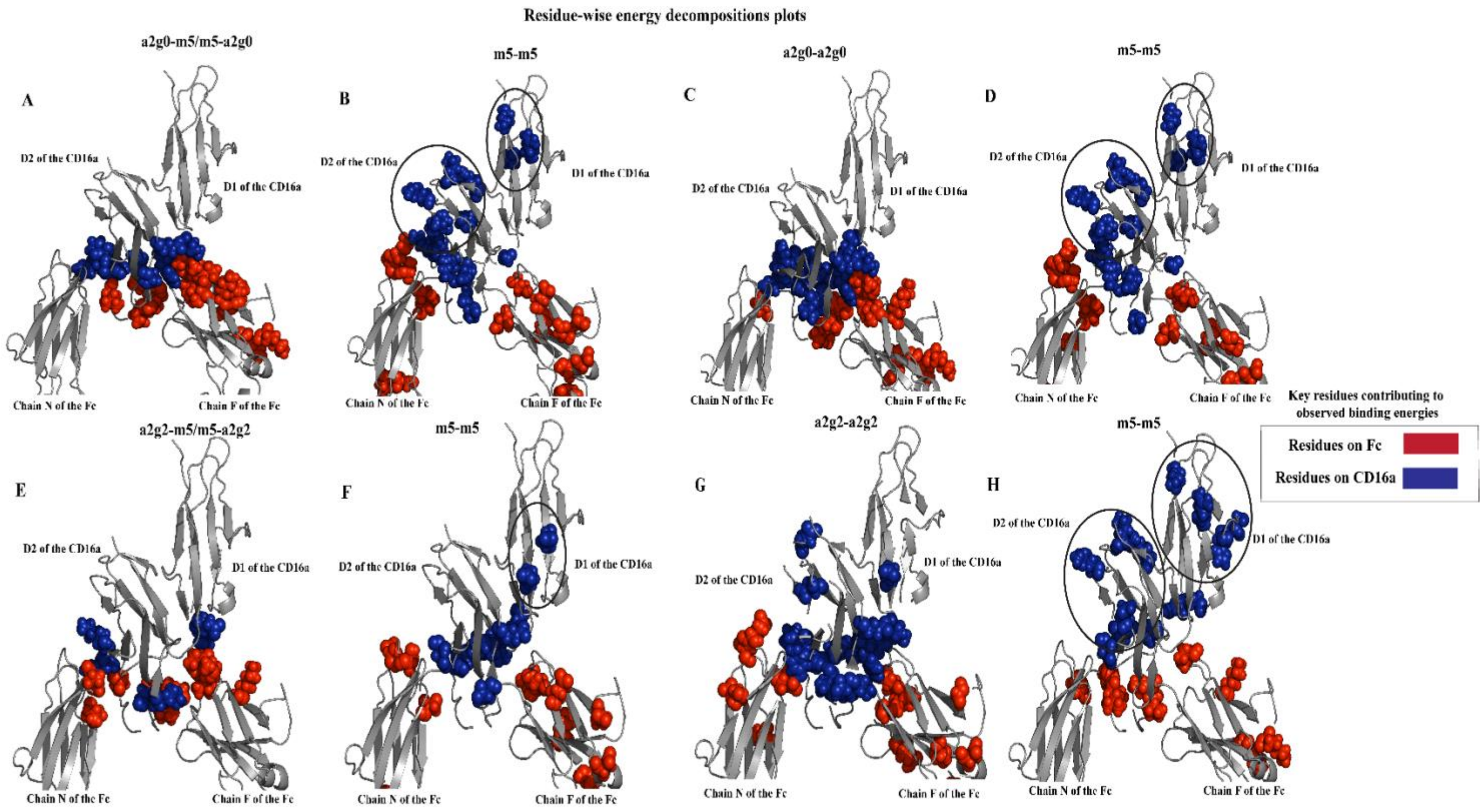


**SI Figure 2: Paired M5 glycosylation fundamentally disrupts the Fc-CD16a binding interface.** Residue-wise energy decomposition analysis identifies the top 20 most energetically active residues at the Fc-CD16a interface for high-affinity Unpaired (G0: **Panel A,** G2: **Panel E**) and Paired biantennary systems (G0: **Panel C,** G2: **Panel G**) and compares them to Paired M5 systems (**Panels B**, **D**, **F** and **H**). Energetically critical CD16a residues in Paired M5 complexes (circled) are substantially displaced from the binding interface relative to both Paired biantennary and Unpaired glycosylated systems, and these observations are in line with **Figure 3**. This spatial redistribution of binding energy indicates that Paired M5 glycosylation fundamentally remodels the interaction landscape and can mechanistically explain the reduced binding affinities observed in **Figure 2**.

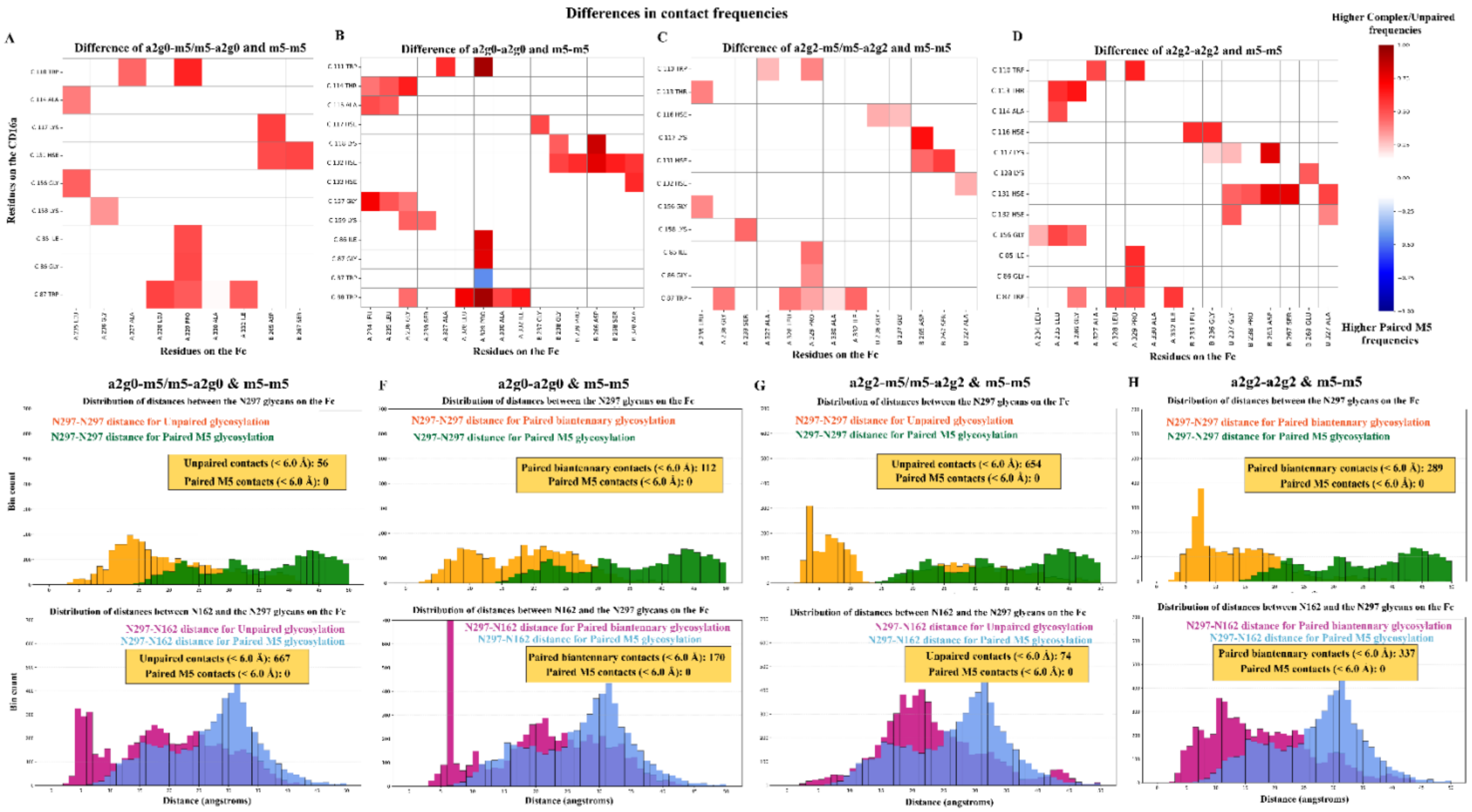


**SI Figure 3: Paired M5 glycosylation minimizes physical contacts at the Fc-CD16a interface.** Contact frequency analysis compares protein-protein (**Panels A**, **B**, **C** and **D**) and glycan-glycan (**Panels E**, **F**, **G** and **H**) interactions between high-affinity Unpaired (**Panels A**, **C**, **E** and **G**) and Paired biantennary systems (**Panels B**, **D**, **F** and **G**) relative to Paired M5 glycosylation. We observe that the Unpaired and Paired systems for G0 (**Panels A** and **B**) and G2 (**Panels C** and **D**) glycosylation exhibit substantially higher protein-protein contact frequencies than the Paired M5 variants, which demonstrates that Paired M5 residues sterically or electrostatically suppress critical inter-protein interactions. Additionally, both Fc N297 inter-glycan contacts and Fc N297–CD16a N162 inter-glycan contacts (**Panels E**, **F**, **G** and **H**) are substantially elevated in Unpaired and Paired biantennary systems compared to Paired M5 glycosylation.

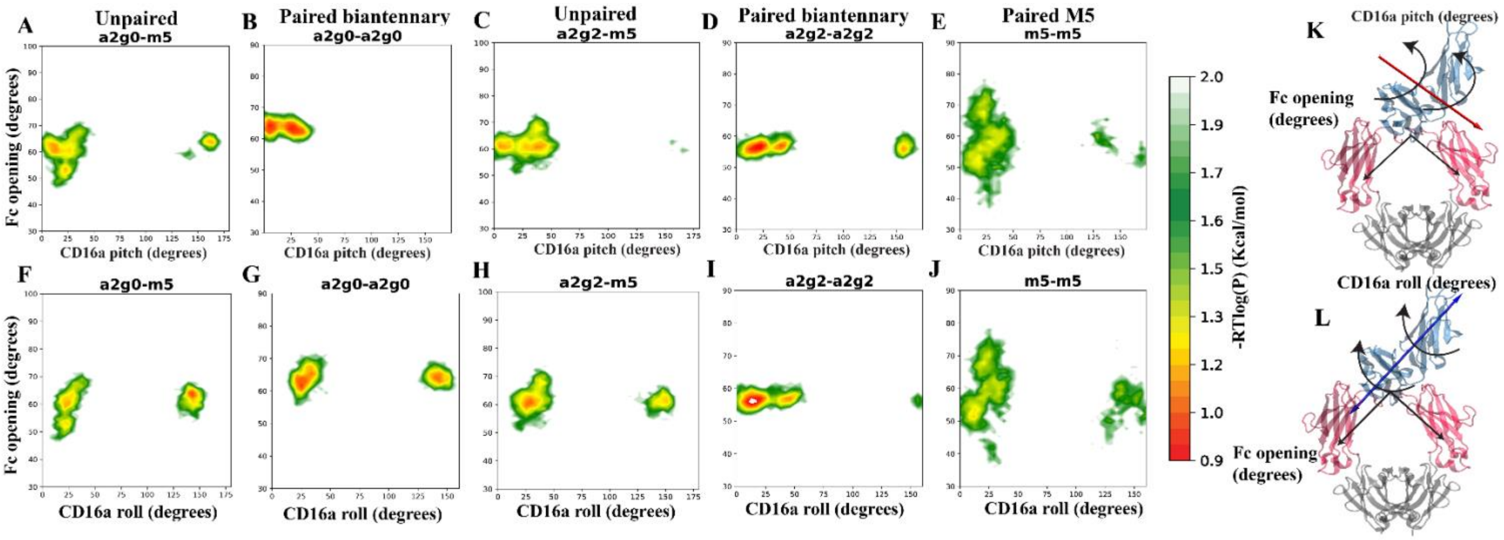


**SI Figure 4: Paired M5 glycosylation destabilizes CD16a conformations and disrupts optimal binding conformations.** Free Energy Surface (FES) analysis describes CD16a rotational motion along two independent axes (**Panels K** and **L**), with lower colorbar values indicating regions of higher stability. Conformational sampling is compared between representative high-affinity Unpaired (G0: **Panels A**, **F** and G2: **C, H**), Paired biantennary (G0: **Panels B**, **G** and G2: **D, I**) systems and the Paired M5 (**Panels E, J**) variants. We find that Paired M5 systems sample substantially larger conformational ensembles with lower stability basins, whereas Unpaired and Paired biantennary systems exhibit discrete, compact energy wells. This conformational destabilization is consistent across both rotational axes, demonstrating that Paired M5 glycosylation fundamentally compromises CD16a conformational stability.

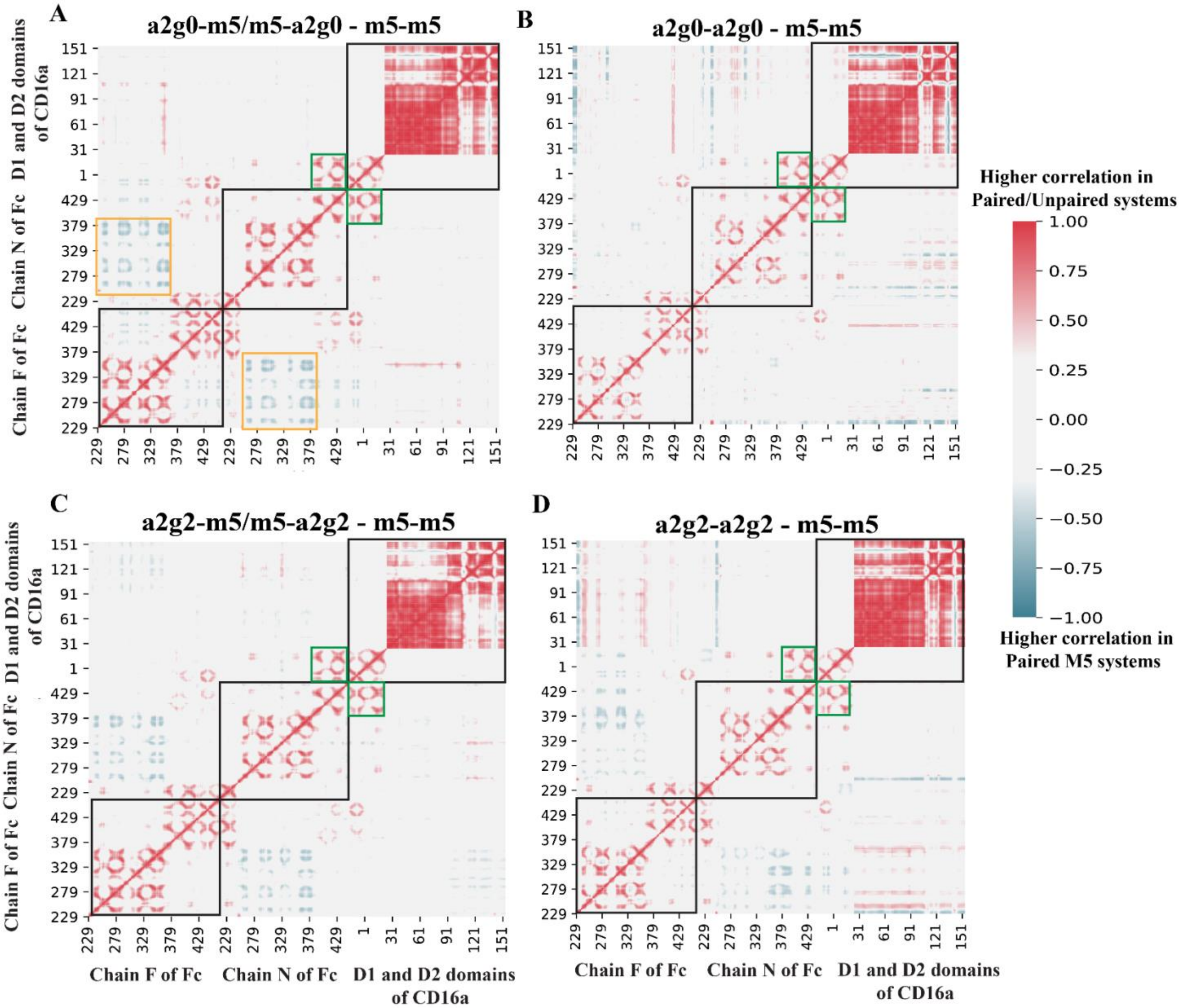


**SI Figure 5: Paired M5 glycosylation leads to a reduction in inter-domain coupling.** Correlation plots describe the differences in correlated motions for Unpaired (**Panels A** and **C**), Paired biantennary (**Panels B** and **D**) systems with respect to Paired M5 systems for G0 (**Panels A** and **B**) and G2 (**Panels C** and **D**) galactosylation levels. We observe higher intra-Fc (see black squares), intra-CD16a (see black squares) and inter Fc-CD16a (see green squares) for both Unpaired and Paired systems, which have been identified to be productive allosteric motions compared to Paired M5 systems.
